# Centromere protection requires strict mitotic inactivation of the Bloom syndrome helicase complex

**DOI:** 10.1101/2024.05.21.595148

**Authors:** María Fernández-Casañas, Eleftheria Karanika, Tomisin Olukoga, Alex Herbert, Umit Aliyaskarova, Matthew Day, Kok-Lung Chan

## Abstract

The BTRR (BLM/TOP3A/RMI1/RMI2) complex resolves various DNA replication and recombination intermediates to suppress genome instability. Alongside PICH, they target mitotic DNA intertwinements, known as ultrafine DNA bridges, facilitating chromosome segregation. Both BLM and PICH undergo transient mitotic hyper-phosphorylation, but the biological significance of this remains elusive. Here, we uncover that during early mitosis, multiple protein kinases act together to strictly constrain the BTRR complex for the protection of centromeres. Mechanistically, CDK1 destabilises the complex and suppresses its association with PICH at the chromatin underneath kinetochores. Inactivating the BLM and TOP3A interaction compromises the UFB-binding complex mitotic functions and can prevent centromere destruction. We further unravel how different clusters of mitotic phosphorylation on BLM affect its interaction with the TOP3A/RMI1/RMI2 subcomplex and illegitimate centromere unwinding. Furthermore, we identify specific phosphorylation sites targeted by the MPS1-PLK1 axis functioning to prevent BLM hyper-activation at centromeres. Notably, unleashing such activity after sister-chromatid cohesion loss facilitates separation of entangled chromosomes. Together, our study defines a centromere protection pathway in human mitotic cells, heavily reliant on a tight spatiotemporal control of the BTRR complex.

## Introduction

Faithful chromosome segregation can be hindered by DNA replication and repair by-products, including double-stranded DNA catenanes, late replication intermediates and homologous recombination structures. These DNA interlinking molecules manifest as nucleosome-poor ultrafine DNA bridges (UFBs) that appear as cells disjoin sister chromatids during anaphase^1–5^. If not resolved properly, they can lead to chromosome mis-segregation, chromatin damage, chromothripsis, and gross chromosomal rearrangements – common characteristic of cancer cells^4–7^. Mammalian cells have evolved a multienzyme complex to specifically target anaphase UFBs, the UFB-binding complex. Its key member is a SNF2-family DNA translocase, PICH (PLK1-interacting checkpoint helicase; also known as ERCC6L)^2^, which recruits the Bloom syndrome DNA helicase, <u>B</u>LM, and its interacting partners <u>T</u>OP3A (Topoisomerase 3A), <u>R</u>MI1 and <u>R</u>MI2 (RecQ-mediated genome instability protein complex 1 and 2; together known as the BTRR dissolvasome)^1,8^. Independently, PICH also recruits RIF1 (RAP1-interacting factor) on anaphase UFBs, but this interaction is suppressed by high levels of CDK1 before anaphase^9^. Apart from these core UFB-binding factors, Polo-like kinase 1 (PLK1) and Protein Phosphatases 1 (PP1s) are also found to associate with PICH and RIF1, respectively, but their biological roles remain unclear. Some components of the UFB-binding complex have important roles during DNA replication and repair, particularly the BTRR dissolvasome is crucial for suppressing excessive sister chromatid exchanges (SCEs)^10–12^. Mutations on BLM and the TRR (TOP3A/RMI1/RMI2) subcomplex have been linked to Bloom syndrome (BS) or BS-like genetic disorders, predisposing individuals to cancer development^13,14^. Given the vital role of these proteins as genome stability guardians, extensive studies have focused on understanding their cellular activities and regulation but mostly in S and G2 cells. Little is known about its role(s) during mitosis, except as core components of the UFB-binding complex.

Upon mitotic entry, both BLM helicase and PICH translocase undergo extensive phosphorylation mediated by several mitotic kinases, including CDK1/2, MPS1 and PLK1^2,15–20^. This is then followed by rapid dephosphorylation at the metaphase-anaphase transition^21^. Over fifty mitosis-specific phosphorylation sites have been reported to occur on BLM, with the majority locating at the N-terminal half of the protein and outside the core helicase domain. The biological functions of most of these phosphorylation events remain unclear, but a relative well studied one is serine 144. It has been suggested that MPS1 phosphorylates BLM at serine 144, creating a docking site for PLK1 binding and subsequent downstream phosphorylation, which is proposed to be involved in facilitating spindle assembly checkpoint (SAC) activation^16^, yet the underlying mechanism remains undetermined. Besides, CDK1 can also phosphorylates BLM at multiple sites^15,18^. Recently, a study has suggested that serine 144 is also a target site of CDK1. In the presence of the binding of TOPBP1 (DNA Topoisomerase 2-binding protein 1) to BLM, it is believed that TOPBP1 can recruit additional PLK1 proteins to accelerate hyperphosphorylation of BLM during the G2-M transition. The lack of the Ser-144 or CDK1-mediated phosphorylation on BLM has been shown to abrogate BLM’s anti-recombination activity in mitosis^18^. This leads to a proposal of mitotic phosphorylation-driven activation of BLM. Interestingly, the phosphorylated form of BTRR complex has been reported to be expelled from mitotic chromosomes^22^ while phosphorylation at threonine 182 of BLM can lead to Fbw7a-dependent polyubiquitylation and protein degradation in mitosis, albeit incomplete^23^. These findings generate a dilemma of how phosphorylated BTRR complexes can effectively access DNA structures within the context of mitotic chromosomes. Nevertheless, a recent study has also suggested that mitotic phosphorylation of BLM is necessary for the assembly and activation of the UFB-binding complex^21^. Whether mitotic cells attempt to activate or deactivate the BTRR complex in mitosis remains controversial, but these studies highly indicate the existence of a regulatory control dependent on the mitotic phosphorylation. Very strikingly, we and others have previously observed a serious mitotic catastrophe, triggered by BLM, when the PLK1 function is inhibited. Rather than the loss of BLM activity, in the absence of PLK1, the BTRR dissolvasome is rapidly recruited to the centromeres by PICH, driving severe centromere dechromatinisation and breakages^20,24^. The centromere damage was found to be initiated via active DNA unwinding by BLM at the chromatin layer underneath the kinetochores^20^. These findings raise several significant questions. Firstly, how does PLK1 protect the centromeres from being attacked by the BTRR dissolvasome? Does it do so by suppressing the BTRR complex activity, and/or by maintaining proper centromere compaction and conformation? Secondly, is the centromere protection reliant on mitotic phosphorylation of the BTRR complex and, if so, what is the underlying mechanism?

In this study, we discover that the BTRR dissolvasome is under a tight, transient inactivation by several mitotic kinases when cells enter mitosis until the onset of anaphase. This is crucial to prevent mis-assembly and activation of the UFB-binding complex at the kinetochore-associated chromatin, which otherwise causes unwanted centromere unwinding and destruction. High levels of CDK1 in early mitosis destabilises the BTRR complex and its binding to PICH, restricting the assembly of the UFB-binding complex. We then identify critical regions and clusters of mitotic phosphorylation on BLM that affect the stability of the BTRR dissolvasome and its DNA unwinding ability at centromeres. Moreover, we find that the MPS1-PLK1 axis mediates specific phosphorylation on BLM to suppress its hyper-activation at centromeric chromatin, whereas the RIF1-PP1s axis is needed to counteract the CDK1’s inhibitory effect during the activation of the UFB-binding complex. Our study thus provides a molecular explanation of why the BTRR dissolvasome needs to be phosphorylated during a specific window of mitosis and, most importantly, it reveals a mechanism operated by a network of mitotic kinases in human cells to safeguard centromere integrity. We generate a comprehensive model to discuss the significance of this centromere protection pathway in the maintenance of chromosome stability.

## Results

### Restriction of the BTRR complex activity before anaphase onset

During normal mitosis, both PICH and BLM are phosphorylated by PLK1^2,16,20^. It is currently unknown whether and how this post-translational modification may contribute to the safeguarding of centromeres. Therefore, we started to investigate the role of PLK1 on regulating the mitotic activity of the BTRR complex. A previous study reported that mitotic phosphorylation can enhance BLM and PICH interaction to promote the formation of UFB-binding complex^21^. However, inhibiting PLK1 with the specific small molecule, BI2536, did not impair the interaction of BLM and PICH, despite leading to the loss of hyper-phosphorylation of both proteins. Instead, it reproducibly enhanced their interaction (Fig. 1A and Supplementary Fig. 1A). Inactivating PLK1 also slightly increased the interaction between BLM and TOP3A (Fig. 1A and Supplementary Fig. 1A). Moreover, although PICH was highly enriched at the <u>k</u>inetochore-associated chromatin (K-chromatin) on metaphase centromeres as shown by high-precision microscopy and super-resolution STED (Supplementary Figs. 1B and 1C), we were unable to detect the binding of the BTRR complex and another PICH-interacting protein, RIF1, unless PLK1 was inhibited (Fig. 1B and Supplementary Fig. 1C). This suggests that the formation of the UFB-binding complex may be negatively regulated before anaphase onset.

**Figure 1.**
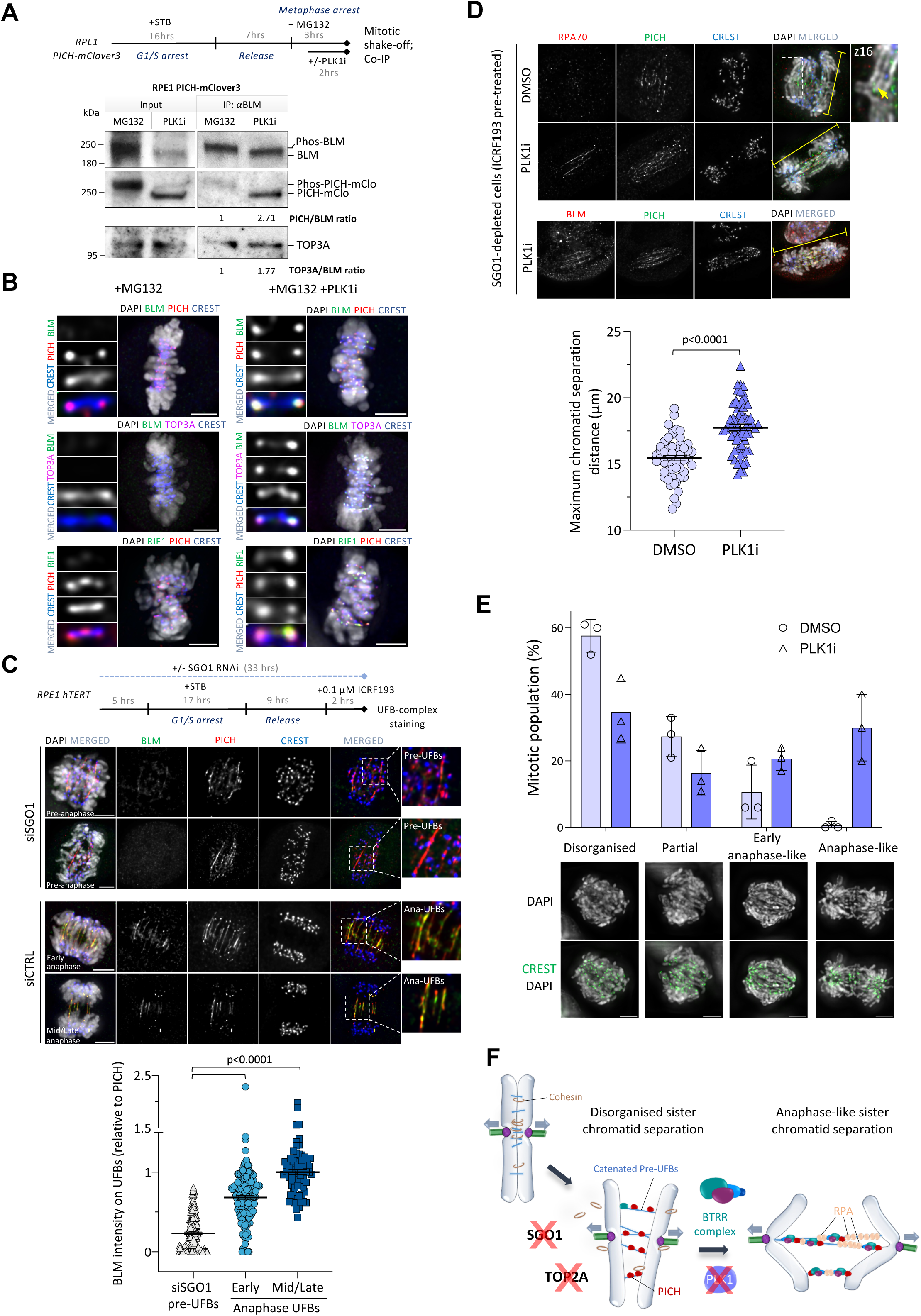
Restricted activation of the UFB-binding complex before anaphase onset. (A) Co-immunoprecipitation analysis in the presence or absence of PLK1i treatments in metaphase-arrested RPE1 cells containing endogenously tagged PICH-mClover3. Endogenous BLM was immunoprecipitated. Quantification of the normalised ratios are shown. (B) The localisation of BLM, TOP3A and RIF1 at K-chromatin in MG132-arrested metaphase cells with or without PLK1i treatments (30 min). RPE1 PICH-mClover3 and HCT116 EGFP-TOP3A cells were stained for BLM; RPE1 EGFP-RIF1 were stained for PICH. Centromeres are labelled by CREST and K-chromatin is marked by PICH. (C) The localisation of BLM on UFBs in RPE1 pre-anaphase and anaphase cells. Experimental outline of SGO1 depletion in RPE1 cells is shown. Representative images of BLM and PICH on pre-anaphase UFBs in SGO1-depleted cells, and on UFBs in the control early and mid/late anaphase cells. BLM was labelled by a rabbit antibody (ab2179). The graph shows the relative intensities (BLM to PICH) on pre-anaphase and anaphase UFB populations. Data is normalised to the average intensity of mid/late anaphase UFBs (total number of UFBs measured: siSGO1 pre-UFBs n=205; early anaphase UFBs n=181; mid/late anaphase UFBs n= 70; mean ± S.E.M is shown). Scale bars, 5 μm. (D) RPE1 cells were treated following the same experimental procedure as (C), but DMSO or PLK1i was added for the last 30 minutes. Representative images of RPA70, BLM and PICH on pre-anaphase UFBs in the SGO1-depleted RPE1 mitotic cells (yellow lines shows the maximum distance of split chromatids). The graph shows the maximum chromatid separation distance (µm) in SGO1-depleted cells pre-treated with ICRF193 in DMSO and PLK1i conditions. (E) Percentages of mitotic populations showing different patterns of sister chromatid separation in the SGO1-depleted RPE1 cells treated with ICRF193 and in the absence or presence of short PLK1 inhibition. Representative images the types of sister chromatid distribution. Scale bars, 5 μm. (F) A cartoon shows the inactivation of PLK1 after premature sister-chromatid cohesion loss promotes anaphase-like separation of catenated chromosomes.

To further examine this phenomenon, we analysed the recruitment of the BTRR complex and RIF1 to UFB structures, but in pre-anaphase cells. RPE1 cells were depleted of SGO1 and treated with a low dose of TOP2 inhibitor, ICRF193, which led to premature loss of sister-chromatid cohesion and the generation of catenated UFB molecules in pre-anaphase cells. We found that both PICH-interacting partners, the BTRR complex and RIF1^9^, poorly localised to the pre-anaphase UFBs that connect split sister chromatids (Fig. 1C and Supplementary Fig. 1D). In contrast, these factors were readily detected along DNA bridges in anaphase, particularly in cells progressing into mid and late anaphase (Fig. 1C and Supplementary Figs. 1D and 1E). In line with the above finding of the induction of K-chromatin localisation, acute inhibition of PLK1 also triggered rapid loading of the BTRR complex onto pre-anaphase UFBs and, crucially, induced DNA bridge unwinding as marked by RPA association (Fig. 1D). Moreover, this led to an increase in the separation of catenated chromosomes (Fig. 1D). Due to the persistent catenation along chromosomal arms (Fig. 1D; arrow), the distribution of the split sister chromatids in the SGO1-depleted cells was generally disorganised and partial. However, this was converted into more anaphase-like patterns after the PLK1i treatments, probably due to the activation of the UFB-binding complex (Figs. 1E and 1F). Therefore, we believe that the activity of BTRR/UFB-binding complex is likely actively suppressed before anaphase by a PLK1-dependent pathway.

### The assembly of a functional UFB-binding complex requires stable BTRR dissolvasome

To understand the underlying mechanism, we decided to determine the PICH-interacting domain(s) on BLM. Following systematic mutagenesis, we identified specific amino acids within the first thirteen residues of BLM that are crucial for proper association with PICH on anaphase UFBs (Fig. 2A and Supplementary Fig. 2A). Since the N-terminal region of BLM has been shown to interact with the TOP3A/RMI1/RMI2 (TRR) subcomplex^25,26^, it is possible that the mutations introduced might disrupt the TRR subcomplex interaction. If this were the case, it would imply that either the TRR subcomplex mediates BLM-PICH interaction, or the formation of BTRR dissolvasome is prerequisite for associating with PICH. However, since Sarlos *et al* have shown that the binding of BLM to PICH does not require the TRR subcomplex^8^, this led us to believe that the very N-terminus of BLM may contain a PICH-interacting motif. Nonetheless, co-immunoprecipitation showed that most of the N-terminal mutants were highly defective in TOP3A pulldown (Fig. 2B). AlphaFold2 structural analyses also predicted that the first fifty residues of BLM, whose structure has not been elucidated, bind across both TOP3A and RMI1 via two consecutive interfaces. The residues from 35 to 50 have been experimentally shown for RMI1 interaction^27,28^ while AlphaFold2 prediction suggested that an α-helix spanning residues 8 to 23 is responsible for TOP3A interaction (Fig. 2C and Supplementary Fig. 2B)^29,30^. Crucially, the point mutations we generated at Pro-5, Asn-8, Leu-9, and Gln-12, which are predicted to stabilise the helix or mediate hydrogen bonding with TOP3A, disrupts BLM-TOP3A interaction (Figs. 2B and 2C).

**Figure 2.**
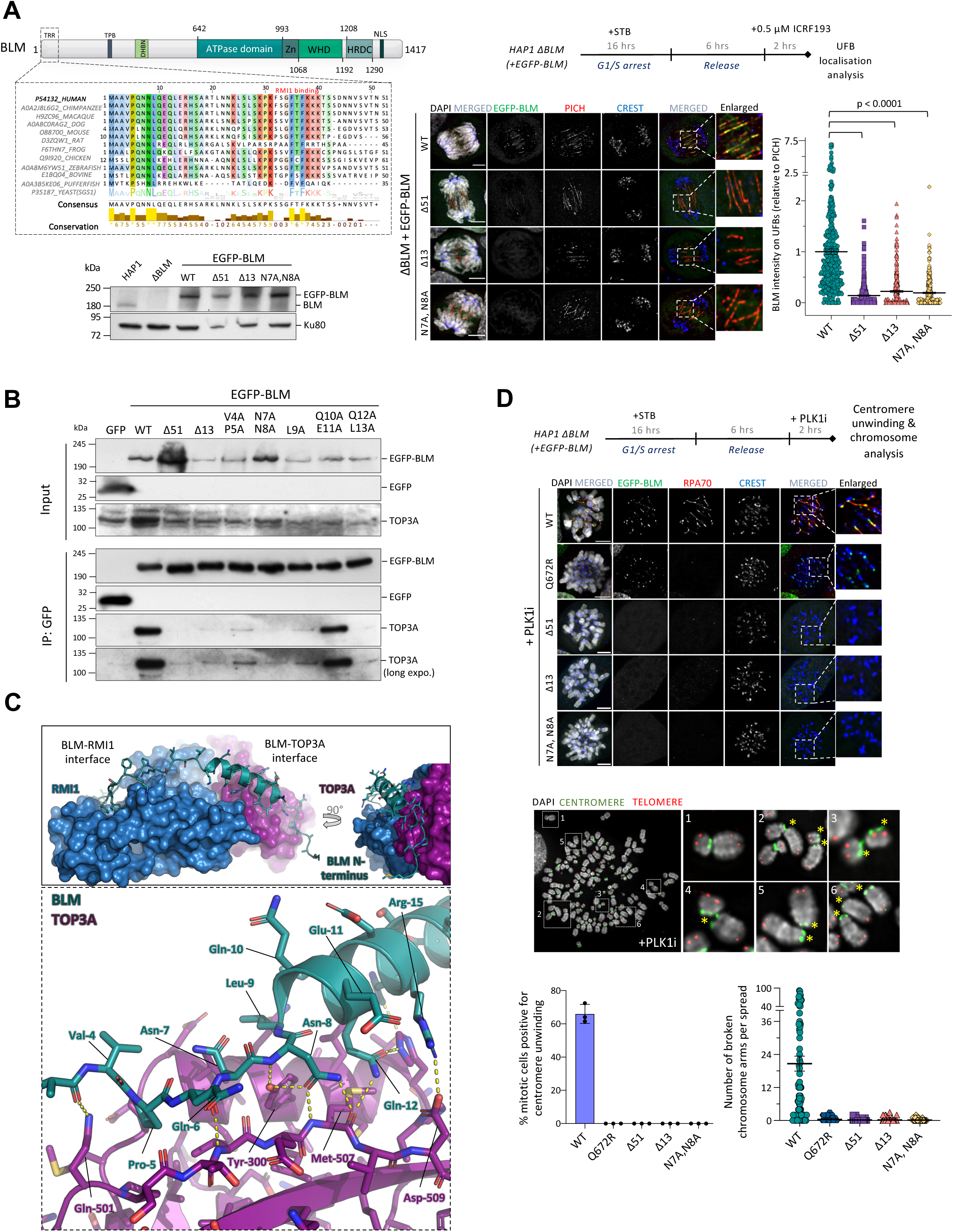
Disruption of BLM and TOP3A interaction impairs UFB-binding complex activities. (A) N-terminal mutations and truncations of BLM impair UFB localisation. Schematic representation shows the sequence alignments and conservation of the first 51 amino acids of human BLM (TRR: TOP3A, RMI1 and RMI2 binding surface; TPB: TOPBP1 binding surface; DHBN; double helical bundle in N-terminus; WHD: Winged Helix domain; HRDC: Helicase and RNaseD C-terminal domain; NLS: Nuclear localisation signal). ΔBLM HAP1 cells were complemented with the indicated EGFP-tagged wildtype and mutants BLM. Western Blot shows the protein levels (Ku80 acts as a loading control). Representative images showing BLM localisation on UFBs labelled by PICH. The graph shows the BLM intensities on UFBs (relative to PICH), normalised to average intensity in WT EGFP-BLM cells (total numbers of UFBs measured: WT=340; Δ51=373, Δ13= 306, N7A, N8A=312; mean ± S.E.M is shown). (B) Mutations of specific N-terminal residues impair TOP3A interaction. EGFP-tagged wildtype and the N-terminal mutants of BLM were transiently expressed in HEK293T cells and subject to co-immunoprecipitation analysis. (C) AlphaFold2 structural prediction model illustrates the N-terminus of BLM as a helix (cartoon representation) with side chains displayed, interacting with both TOP3A and RMI1 (surfaces) through two distinct interfaces. The interacting residues are highlighted and shown as sticks, with hydrogen bonds represented as dashed lines. (D) N-terminal mutations of BLM impair K-chromatin binding and centromere unwinding induced by PLK1 inhibition. (Top) Representative images of ΔBLM HAP1 mitotic cells complemented with EGFP-tagged wildtype, helicase-dead (Q672R), and N-terminal mutant BLM after PLK1i treatment (100 nM BI2536). (Middle) Representative images of metaphase spread chromosomes showing centromere breakages induced by PLK1i treatments (60 nM BI2536) (asterisks). Chromosomes were hybridised with FISH DNA probes against centromeres (green) and telomeres (red). The left graph shows percentages of mitotic cells positive for centromere unwinding (total number of cells analysed: WT n=269; Q672R n= 182; Δ51 n=179; Δ13 n=183; N7A, N8A n=184; mean ± S.D. is shown, three independent experiments). Scale bars, 5 μm. The right graph shows quantification of the number of broken whole chromosome arms per spread after PLK1i treatments in ΔBLM HAP1 complemented cells with EGFP-tagged BLM wildtype and mutants (Q672R and N-terminal mutants: Δ51, Δ13 and N7A, N8A; at least 80 spreads were analysed per condition; mean ± S.E.M is shown).

To strength the evidence that this is a *bona fide* TOP3A interacting motif, we set up a cellular protein-protein interaction assay. EGFP-tagged wildtype, helicase-dead, and the N-terminal mutants of BLM were tethered to centromeres by fusion to a CENPB peptide, which served as a “bait” to investigate TOP3A interaction *in vivo* (Supplementary Fig. 2C). Both the wildtype and helicase-dead BLM fusion proteins effectively recruited TOP3A to centromeres, indicating the presence of BLM-TOP3A interaction and that is independent of the helicase activity. Mutating the putative TOP3A-binding motif completely abolished the centromeric recruitment of TOP3A (Supplementary Fig. 2C). Collectively, this demonstrates that the binding of BLM to PICH on UFBs requires TOP3A interaction. Notably, we found that the TOP3A-binding mutants were also defective in K-chromatin localisation, and failed to induce centromere unwinding and breakages following PLK1 inhibition^20^ (Fig. 2D). These data indicate that the mitotic activity of the BTRR complex requires its stable formation, which is inconsistent with the previously proposed model^8^. To corroborate our findings, we depleted TOP3A in different cell lines, which also abolished BLM localisation to UFBs (Supplementary Fig. 3A). Likewise, the ablation of BLM also prevented TOP3A from binding to PICH on UFBs (Supplementary Figs. 3B and 3C). Therefore, we conclude that the assembly and activity of the UFB-binding complex in mitosis relies on intact BTRR complex.

### Disassembly of the mitotic BTRR dissolvasome at centromeres

Given this unexpected finding, we hypothesised that one potential mechanism to protect centromeres against the attack by the BTRR/UFB-binding complexes during mitosis could involve destabilisation of the BTRR complex. To test this possibility, we combined the cellular protein-protein interaction assay and real-time microscopy to determine the interaction dynamics between different subunits of the BTRR complex in live cells. We tethered either mCherry-tagged RMI1 or helicase-dead BLM(Q672R) proteins to centromeres through CENPB fusion and monitored the recruitment of EGFP-TOP3A protein throughout cell division (Fig. 3A). A helicase-dead BLM(Q672R) was used to minimise potential interferences to centromeric structures. As planned, the RMI1 and BLM(Q672R) fusion proteins were successfully immobilised at centromeres in both interphase and mitotic cells (Fig. 3B). In the CENPB-mCherry-RMI1 expressing cells, EGFP-TOP3A co-localised with RMI1 at the centromeres from G2 to M phases, and throughout mitosis. There was no obvious change in the TOP3A recruitment efficiency (Figure 3B, upper panel; Movie S1), indicating that the TOP3A-RMI1 interaction is constantly maintained, regardless of the cell cycle stage. Intriguingly, we detected a reduction of TOP3A localisation to the centromeres in CENPB-mCherry-BLM(Q672R) cells upon mitotic entry. Remarkably, the BLM-TOP3A interaction was quickly restored after anaphase onset (Figure 3B, lower panel; Movie S2). Similar changes of the interaction dynamics between endogenous BLM and the CENPB-mCherry-RMI1 proteins were also detected by quantitative immunofluorescence microscopy (Figure 3C). These data support our prediction that the BTRR complex can be destabilised, especially during early mitosis. However, in soluble protein extracts of mitotic cells, the formation of BTRR complexes has been detected^22,31^. Our *in vivo* protein-protein interaction analysis thus may indicate that the stability of the complex is susceptible to centromere chromatin environment and/or the complex becomes intrinsically less stable in mitosis. We further examined this by subjecting the BTRR complex immunoprecipitated from interphase and mitotic cell extracts to different salt conditions. Repeatedly, we detected decreased pull-down efficiencies between BLM and TOP3A in mitotic extracts from different cell lines (Supplementary Figs. 4A and 4B). The unstable nature of the mitotic BTRR complex at centromeres may hinder its proper association with PICH at the K-chromatin, thereby preventing mis-activation of the UFB-binding complex.

**Figure 3.**
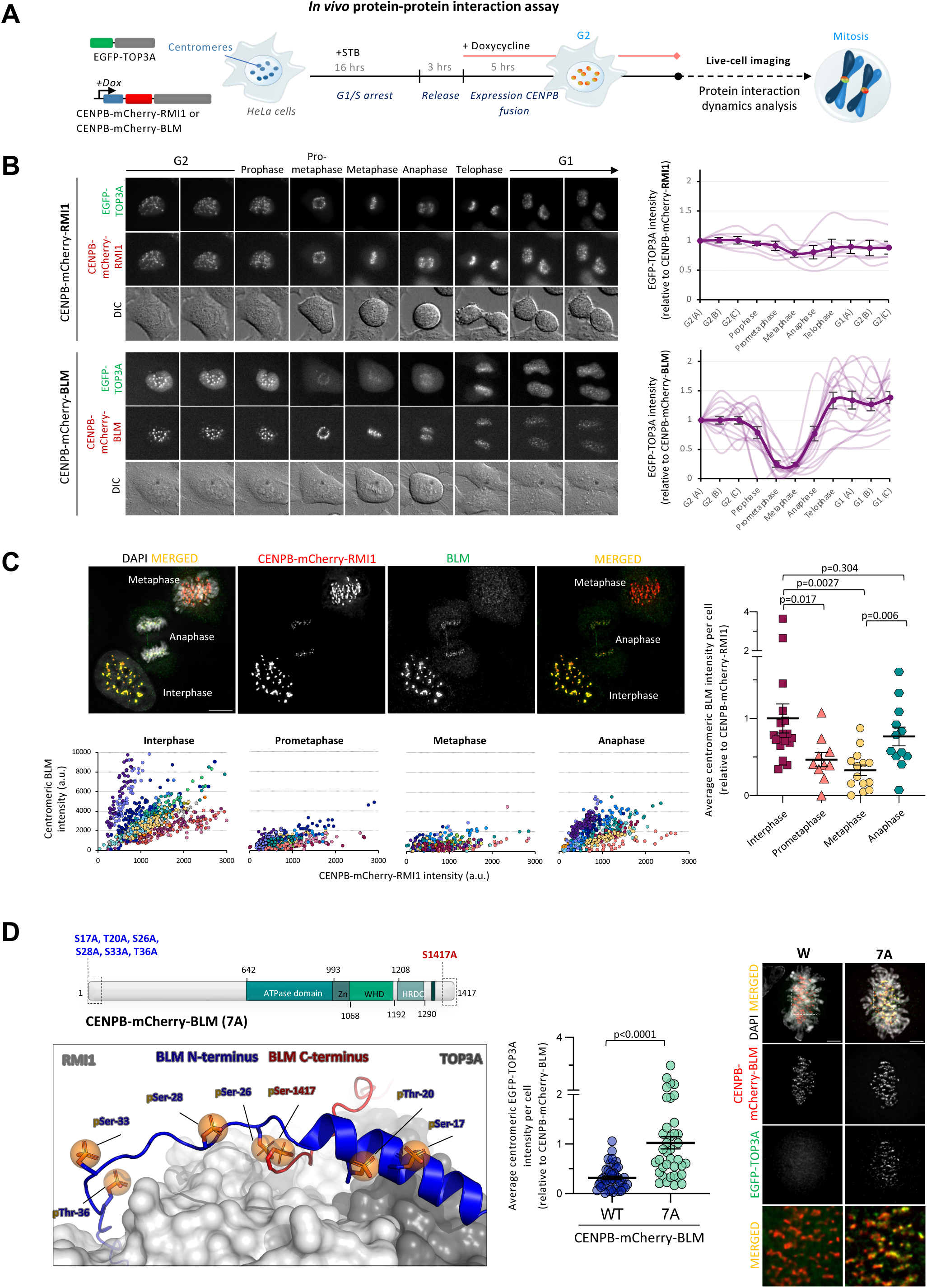
Dynamic changes of the BLM-TOP3A interaction at centromeres during mitosis. (A) Diagram showing an *in vivo* live-cell imaging assay to measure the dynamic changes of protein-protein interaction at centromeres in HeLa cells expressing EGFP-TOP3A, and either CENPB-mCherry-RM1 or -BLM fusion proteins. (B) Time-lapse live-cell images showing the TOP3A-RMI1 interaction (upper panel) and TOP3A-BLM interaction (lower panel) at centromeres from late G2 to the next G1 phase. Graphs show quantification of centromeric EGFP-TOP3A intensities relative to CENPB-mCherry-RMI1 or -BLM in the indicated cell-cycle stages (number of cells analysed: CENPB-mCherry-RMI1 n=7; CENPB-mCherry-BLM n=16; mean ± S.E.M is shown). (B) Quantitative immunofluorescence microscopy showing the recruitment of endogenous BLM proteins to centromeres by the CENPB-mCherry-RMI1 in HeLa interphase and mitotic cells. Scales bars, 10 µm. Scatter plots showing the intensities of BLM versus CENPB-mCherry-RMI1 at individual centromere clusters (each colour represents different cells examined; numbers of centromere clusters measured: Interphase n=933; Prometaphase n= 336; Metaphase n=411, Anaphase n=616). The right graph shows quantification of average centromeric BLM intensity relative to CENPB-mCherry-RMI1 per cell. Data is normalised to the average intensity in metaphase (numbers of cells analysed: Interphase n=19; Prometaphase n= 10; Metaphase n=14, Anaphase n=12; mean ± S.E.M is shown). (C) A diagram shows the mitotic phosphorylation sites on BLM in the N- and C-termini that span the interacting surfaces for TOP3A and RMI1 (See also Fig. S2). AlphaFold2 model illustrates the BLM as a helix in cartoon representation rainbow coloured from N-terminus (blue) to C-terminus (red), while TOP3A and RMI1 are depicted as surfaces. Phosphorylated residues on BLM are represented as spheres. Immunofluorescence images showing the co-localisation of EGFP-TOP3A and CENPB-mCherry-BLM(7A) mutant at centromeres in interphase and metaphase HeLa cells. Immunofluorescence images comparing the centromeric recruitment of EGFP-TOP3A by CENPB-mCherry-BLM(Q672R) and CENPB-mCherry-BLM(7A) in metaphase cells. The graph shows quantification of average centromeric EGFP-TOP3A intensity relative to CENPB-mCherry-BLM per cell. (numbers of cells analysed: CENPB-mCherry-BLM n=42; CENPB-mCherry-BLM 7A n= 37; mean ± S.E.M is shown).

### Mitotic phosphorylation weakens the BLM and the TRR-subcomplex interacting surfaces

Several mitotic kinases accumulate at the centromere, and they have been shown to phosphorylate the subunits of the BTRR complex^15,16,19,20^. We next tested whether they may involve in destabilising the complex during early mitosis. The CENPB-mCherry-RMI1 metaphase-arrested cells were treated with different specific inhibitors targeting PLK1, CDK1, MPS1, Aurora A or Aurora B. A short treatment time was applied to avoid triggering premature mitotic exit, particularly during CDK1 and MPS1 inhibition^32–34^. We found that inhibition of PLK1 or CDK1 could effectively restore the binding of BLM to the centromere-tethered RMI1, while Aurora B inhibition exhibited a partial effect (Supplementary Fig. 4C). By integrating data obtained from mass spectrometry analyses and AlphaFold2 prediction studies, we identified six potential mitotic phosphorylation sites on the N-terminus of BLM, including Ser-17, Thr-20 in the helical region, and Ser-26, Ser-28, Ser-33, and Thr-36, as well as one at the C-terminus, Ser-1417^29,35–38^. Potentially, phosphorylation of these sites might disrupt the structural integrity of the TOP3A- and RMI1-binding interfaces, affecting the interaction between BLM and the TRR subcomplex (Fig. 3D). As predicted, mutations at these seven residues, CENPB-mCherry-BLM (7A), rescued the loss of EGFP-TOP3A centromeric recruitment in early mitotic cells (Fig. 3D). This indicates a phosphorylation-dependent mechanism to destabilise the BTRR complex at centromeres.

### CDK1 and MPS1-PLK1 differentially govern the assembly and activity of the UFB-binding complex

Next, we tested whether stabilising the mitotic BTRR complex was able to override the suppression on binding to PICH at K-chromatin. EGFP-BLM expressing HAP1 ΔBLM cells and RPE1 cells were arrested in metaphase and treated with different kinase inhibitors (Fig. 4A). PLK1 inhibition, as expected, effectively induced the loading of BTRR dissolvasome onto K-chromatin. However, we found that only inhibition of CDK1, but not Aurora B, can induce the K-chromatin binding, despite both being able to restore the mitotic BTRR complex formation at centromeres (Fig. 4B and Supplementary Fig. 5A). This indicates that PLK1 and CDK1 also block the complete BTRR complex from binding to PICH. More intriguingly, we noticed that although the BTRR complexes accumulated at K-chromatin following CDK1 inhibition, centromeric DNA unwinding was poorly detected, unlike under the PLK1i treatments (Fig. 4C). This data strongly implies that PLK1 has an additional function to suppress BLM mitotic activity and/or protect the centromeric structures from being recognised as a BLM DNA substrate.

**Figure 4.**
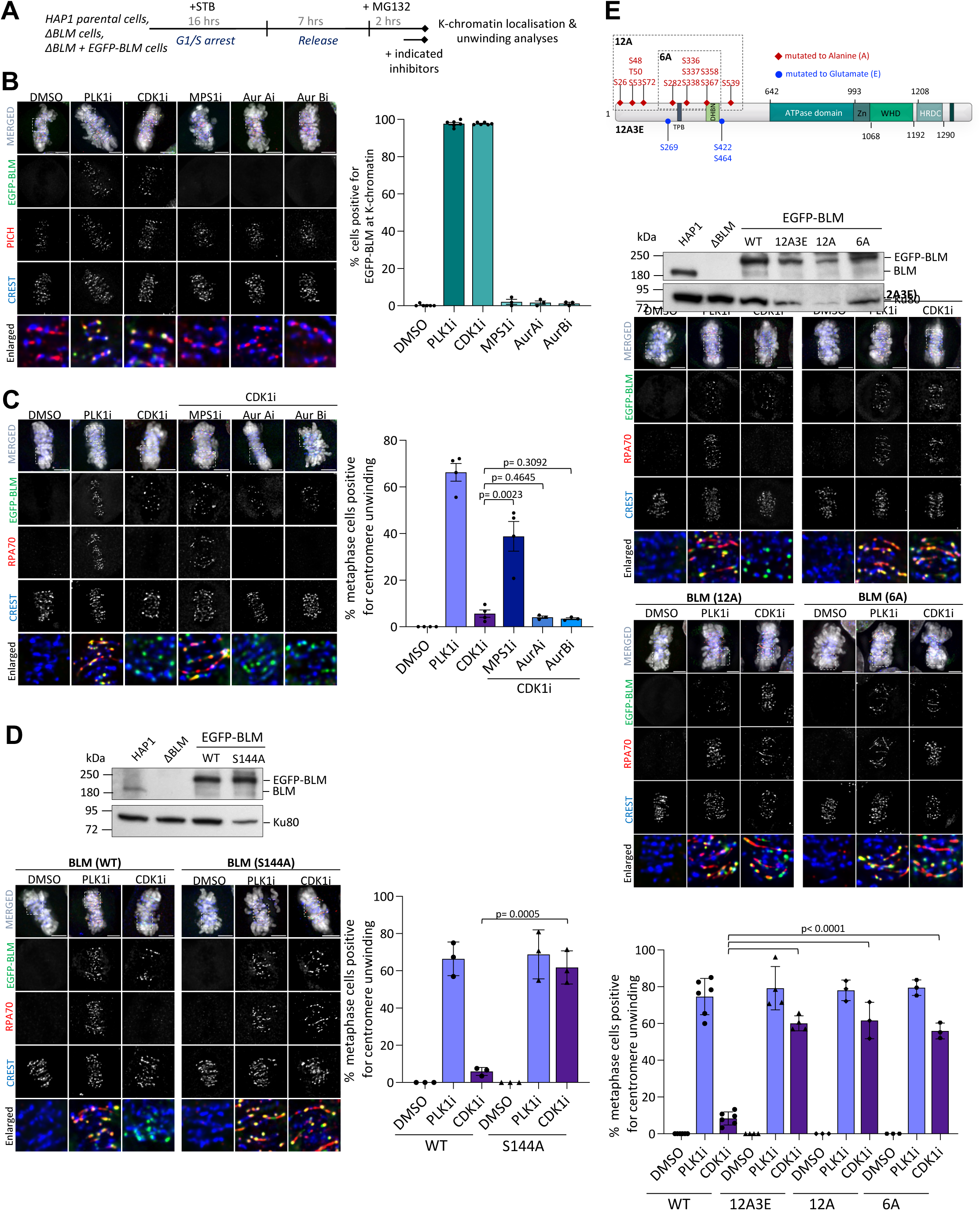
CDK1 and MPS1 kinases differentially regulate the assembly and activity of the UFB-binding complex. (A) Experimental procedures of K-chromatin localisation and centromere unwinding analyses. (B) CDK1 or PLK1 inhibition induces BLM localisation to the K-chromatin. Representative images of ΔBLM HAP1 cells expressing EGFP-BLM arrested in metaphase followed by the indicated kinase inhibitor treatments. PICH labels for K-chromatin and CREST for centromeres. Quantification of the percentage of cells positive for BLM at K-chromatin (number of metaphase cells analysed: DMSO n=175; PLK1i n= 201; CDK1i n=210; MPS1i n=170; AurAi n=177; AurBi n=154; mean ± S.D. is shown, from at least three independent experiments). (C) Similar to (B) but containing co-treatments of the indicated inhibitors. Images and quantification of percentages of cells positive for centromere DNA unwinding - the presence of RPA threads or foci (total numbers of metaphase cells analysed: DMSO n=243; PLK1i n= 247; CDK1i n=268; CDK1i/MPS1i n=200; CDK1i/AurAi n=213; CDK1i/AurBi n=227; mean ± S.D. is shown from three independent experiments). (D) Western Blot shows the EGFP-BLM protein levels in the ΔBLM HAP1 cells. Representative images of ΔBLM HAP1 cells expressing EGFP-tagged wildtype or S144A phosphomutant BLM after PLK1i or CDK1i treatments. Quantification of percentages of metaphase cells positive for centromere DNA unwinding (total numbers of metaphase cells analysed in EGFP-BLM WT: DMSO n=197; PLK1i n= 205; CDK1i n=224 and EGFP-BLM S144A: DMSO n=165; PLK1i n= 183; CDK1i n=188; mean ± S.D. is shown, from three independent experiments). (E) Schematic representation showing serine/threonine residues mutated to alanine or glutamate in the EGFP-BLM phosphomutants; 12A3E (S26A, S48A, T50A, S53A, S72A, S282A, S336A, S337A, S338A, S358A, S367A, S539A, S269E, S422E, S464E); 12A (similar to 12A3E but excluding the three glutamate substitutions), 6A (S282A, S336A, S337A, S338A, S358A, S367A). Western Blot shows the EGFP-BLM protein levels in the ΔBLM HAP1 cells complemented with EGFP-BLM 12A3E, 12A and 6A. Representative images showing EGFP-BLM and RPA foci at K-chromatin in wildtype, 12A3E, 12A or 6A phosphomutants of BLM after the indicated treatments. Quantification of the percentages of metaphase cells positive for centromere DNA unwinding (total numbers of metaphase cells analysed in EGFP-BLM WT: DMSO n=348; PLK1i n= 431; CDK1i n=403, EGFP-BLM 12A3E: DMSO n=237; PLK1i n= 382; CDK1i n=363, EGFP-BLM 12A: DMSO n=209; PLK1i n= 225; CDK1i n=289, EGFP-BLM 6A: DMSO n=201; PLK1i n= 218; CDK1i n=263; mean ± S.D. is shown, from at least three independent experiments). Scale bars, 5 µm. See also Supplementary Fig. 5.

To dissect these possibilities, we studied the function of other mitotic phosphorylation on BLM. It has been suggested that MPS1 catalyses a priming phosphorylation on BLM at serine 144, leading to downstream PLK1-mediated hyper-phosphorylation^16^. If BLM mitotic activity could be directly inhibited by this MPS1-PLK1 axis, we suspected that inactivating MPS1 could also bypass the suppression of centromere DNA unwinding once the BTRR complex gained the access to K-chromatin via CDK1 inhibition. As predicted, acute co-treatment of both CDK1 and MPS1 inhibitors, mimicking PLK1 inhibition, successfully induced single-stranded DNA formation at centromeres, albeit slightly milder (Fig. 4C). To exclude the possibility that this is a secondary effect of MPS1 activity loss, we stably expressed wildtype and S144A phosphomutant BLM in ΔBLM cells. Consistent with our hypothesis, cells containing the S144A BLM displayed a six-fold increase in centromere DNA unwinding activity as compared to the wildtype counterpart (Fig. 4D). Therefore, MPS1-dependent Ser-144 phosphorylation can suppress mitotic BLM activity at centromeres. Our data is somewhat inconsistent with a recent report claiming that the lack of Ser-144 phosphorylation impairs BLM activity such as sister chromatids exchange (SCE) suppression^18^. We thus measured SCE frequencies in the S144A mutant cells but did not detect any obvious defects (Supplementary Fig. 5B). Therefore, we believe that Ser-144 phosphorylation functions to limit, rather than stimulate, BLM activity during mitosis.

To further test whether the suppression of centromere unwinding is dependent of the downstream PLK1 phosphorylation, we performed mass spectrometry analyses and comprehensive phosphoproteomic comparison^35,37,39–43^, and identified twelve phosphorylated serine/threonine sites on BLM that are highly dependent on PLK1 activity. Interestingly, three phosphorylation events were also observed after PLK1 inactivation (Supplementary Fig. 5C). We then complemented ΔBLM cells with phosphomutants of BLM containing the corresponding 12A mutations with or without additional three phospho-mimicking glutamate substitutions (12A3E or 12A) (Fig. 4E). Consistent with the finding from the S144A mutant cells, both 12A and 12A3E BLM mutants also exhibited a significant elevation of K-chromatin unwinding following CDK1 inhibition. Further mutagenesis revealed a small cluster of PLK1-dependent phosphorylation sites (6A) locating in the middle of the flexible N-terminal region of BLM crucial for suppressing the illegitimate centromere unwinding (Fig. 4E). Taken together, we conclude that BLM mitotic activity at the centromere is directly suppressed by a cluster of phosphorylation mediated by PLK1 spanning in the middle of the N-terminal region of BLM (Supplementary Fig. 5D). In parallel, CDK1 and PLK1 can destabilise the BTRR dissolvasome and also prevent it from binding to PICH on K-chromatin. These combined actions create a robust two-tier inhibitory system protecting the centromeres from the unwanted activity of the UFB-binding complex.

### RIF1-PP1s are an auxiliary factor for the UFB-binding complex activation before anaphase

Our findings thus far strongly indicate that PLK1 acts as a major suppressor of the UFB-binding complex during mitosis. However, unexpectedly, we found that inhibiting PLK1 in cells lacking RIF1 poorly induced the loading of the BTRR complex to K-chromatin and centromere unwinding, albeit not completely absence (Fig. 5A). This was not due to the retention of PLK1-dependent hyper-phosphorylation (Fig. 5B). However, inhibiting CDK1 in the ΔRIF1 cells was still effective to induce the UFB-binding complex assembly on K-chromatin (Fig. 5C). Therefore, this implies that CDK1, rather than PLK1, is primarily responsible for blocking the UFB-binding complex formation. If this was the case, how do RIF1-proficient cells overcome the CDK1 inhibitory function? RIF1 has been shown to interact and recruit Protein Phosphatases 1 (PP1s) to PICH^21^. We hypothesised that during the PLK1 inactivation, the RIF1-PP1s interaction and/or PP1s activity were restored, which could counteract the CDK1-dependent suppression. As such, the binding between RIF1 and PP1s, coupled with PP1s activity, is likely to be critical. Supporting this notion, we found that centromere unwinding and breakages were inefficiently induced in RPE1 ΔRIF1 cells complemented with a PP1-binding mutant of RIF1 (PP1bs)^44^ (Figs. 5D and 5E). Moreover, treating cells with a specific inhibitor, tautomycetin^45^, targeting PP1s enzymatic activity also hindered the binding of the BTRR dissolvasome to K-chromatin induced by PLK1 inhibition (Fig. 5F). Collectively, these results demonstrate that the regulation of spatiotemporal activity of the BTRR and UFB-binding complexes is governed by different actions exerted by both mitotic kinases and phosphatases in order to prevent accidental damage to the centromeres.

**Figure 5.**
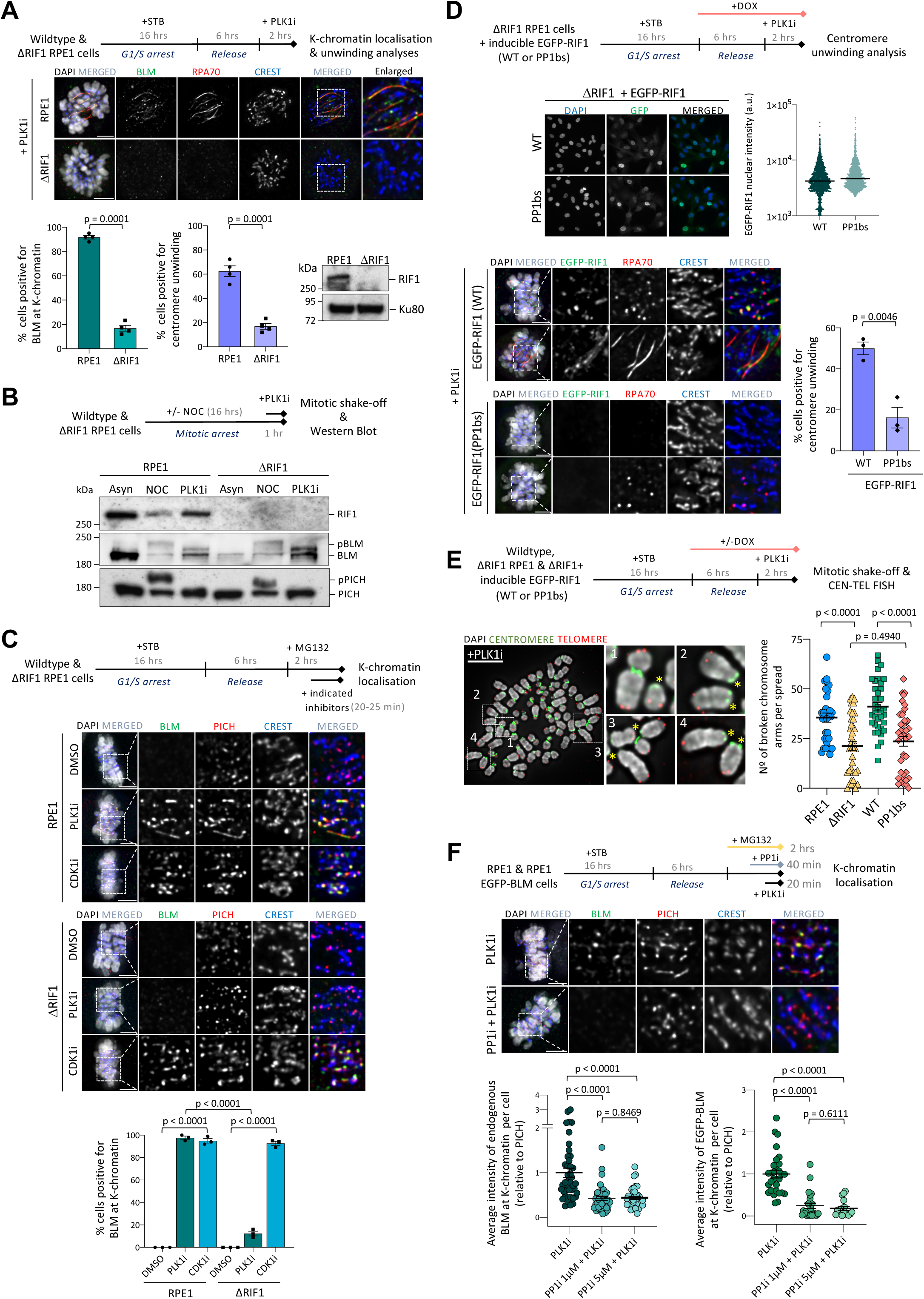
RIF1 and PP1s facilitate the activation of UFB-binding complex. (A) K-chromatin localisation and centromere unwinding in wildtype and ΔRIF1 RPE1 cells treated with PLK1i. Representative images and graphs of the percentages of mitotic cells positive for BLM localisation at K-chromatin (left) and centromere unwinding (right) are shown (numbers of cells analysed: RPE1 n=282; ΔRIF1 n= 295; mean ± S.D. is shown, from four independent experiments). RIF1 expression was analysed by Western Blotting. (B) Electrophoretic mobility shifts of BLM and PICH following PLK1i treatments in wildtype and ΔRIF1 RPE1 cells. (C) K-chromatin localisation of BLM after the indicated treatments in wildtype and ΔRIF1 RPE1 cells. Experimental outline, representative images, and graph of percentages of metaphase-arrested cells positive for BLM at K-chromatin are shown (numbers of cells analysed in RPE1: DMSO n=184; PLK1i n= 195; CDK1i n=198 and in ΔRIF1: DMSO n=175; PLK1i n= 202; CDK1i n=210; mean ± S.D. is shown, from three independent experiments). (D) Centromere unwinding analysis in ΔRIF1 RPE1 cells with Dox-inducible wildtype (WT) or PP1s-binding mutant (PP1bs) of EGFP-RIF1. RIF1 expression was measured by quantitative immunofluorescence staining. Representative images and quantification of the percentage of mitotic cells positive for centromere unwinding in WT and PP1s-binding mutant RIF1 cells are shown (total numbers of cells analysed in WT n=218 and in PP1bs n=234; mean ± S.D. is shown, from three independent experiments). Scale bars, 5 µm. (E) Centromere disintegration analysis. Representative images of metaphase chromosomes after PLK1 inhibition. Centromeres and telomeres are labelled by FISH PNA probes. Asterisks indicate chromatin with centromere breakage. Quantification of broken whole chromosome arms per metaphase spread (number of metaphase spreads analysed: RPE1 n=31; ΔRIF1, n=39; ΔRIF1 EGFP-RIF1 WT n= 37; ΔRIF1 EGFP-RIF1 PP1bs n= 42. mean ± S.E.M is shown). (F) Experimental outline of co-treatments of PP1s and PLK1 inhibitors. Representative images of BLM K-chromatin localisation in RPE1 metaphase cells. Graphs show quantification of average centromeric intensity of BLM relative to PICH per cell in RPE1 (left) and RPE1 EGFP-BLM (right) cells. Data is normalised to the average intensity in PLK1i condition (number of cells analysed in each condition: RPE1 cells at least 40; RPE1 EGFP-BLM at least 20; mean ± S.E.M is shown).

## Discussion

Considering the brief duration of anaphase, it is reasonable to believe that the UFB-binding complex factors may have been activated prior to anaphase, which, in principle, can facilitate rapid separation of sister chromatids thereafter^18,21^. Contrary to this concept, our current study reveals that human cells actively constrain the assembly and spatiotemporal activity of the UFB-binding complex before chromosome segregation. This suppression is vital to prevent illegitimate DNA unwinding at the kinetochore-associated chromatin (K-chromatin), an initiation modification that leads to centromere breaks and hence failure of chromosome alignment^20,24^. To avoid such mitotic catastrophe, we show that high levels of CDK1 in mitotic cells can destabilise the BTRR dissolvasome and prevent its association with PICH at K-chromatin, thus blocking access to the centromeres. Concurrently, MPS1 and PLK1 kinases collaborate to phosphorylate BLM at specific residues, which limits its DNA unwinding ability at the centromeres (Fig. 6A). This two-tier inhibitory system offers several advantages. Firstly, it prevents the BTRR complex from interfering with chromatin regions where only PICH activity is required, such as at the K-chromatin and along chromosomal axes^46^. Secondly, the suppression of BLM activity minimises unwanted chromatin remodelling, even if the complex is misplaced. Additionally, because PICH can still associate with UFB precursors, rapid assembly of the UFB-binding complex can be achieved upon the removal of the inhibitory phosphorylation at the metaphase-anaphase transition. This swift process is highly beneficial for timely segregation of chromosomes. Indeed, we have demonstrated that acute PLK1 inhibition, following the inactivation of sister chromatid cohesion, can promote the separation of entangled chromosomes, presumably through rapid unwinding of the DNA bridges.

**Figure 6.**
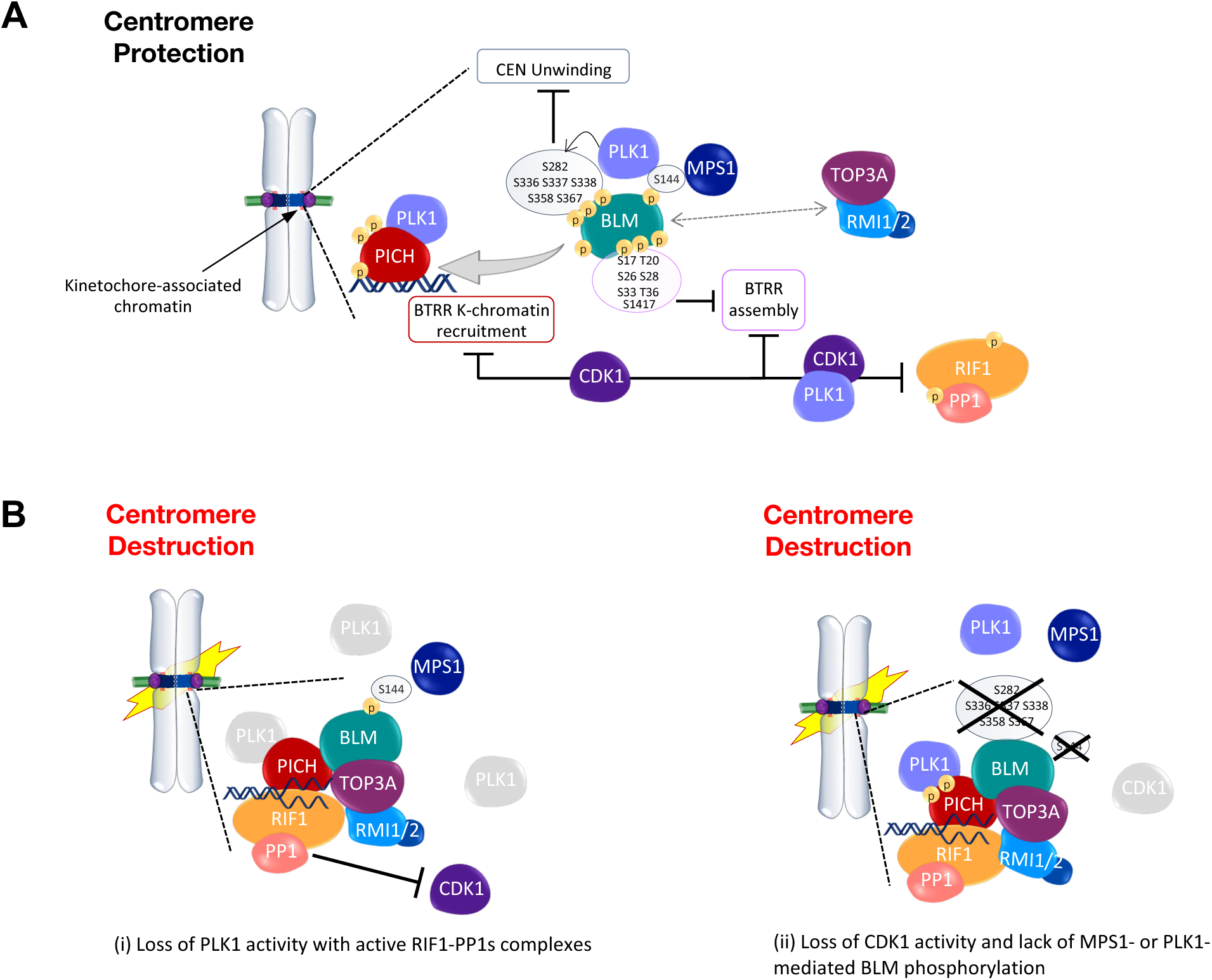
A model of the centromere protection pathway. (A) Multiple mitotic kinases work cooperatively to prevent the activation of the UFB-binding complex before anaphase onset. They suppress stable formation of the BTRR complex and its association with PICH at K-chromatin as well as the DNA unwinding reaction at centromeres. CDK1 can block stable formation of the BTRR dissolvasome and UFB-binding complexes. Mitotic phosphorylation of BLM at Ser-17, Thr-20, Ser-26, Ser-28, Ser-33, Ser-36 and Ser-1417 reduces the binding to the TOP3A/RMI1/RMI2 subcomplex at centromeres. Disrupting the BLM-TOP3A interaction destabilises the BTRR complex assembly and the binding to PICH at K-chromatin and on UFB structures. MPS1-dependent phosphorylation on Ser-144 promotes PLK1 binding, leading to further BLM phosphorylation at residues including Ser-282, Ser-336, Ser-337, Ser-338, Ser-358 and Ser-367, which can suppress centromeric DNA unwinding by the BTRR complex. (B) Dysregulation of the mitotic inhibitory system causes K-chromatin DNA unwinding and centromere destruction. This can be caused by (i) the loss of PLK1 activity, which leads to the activation of RIF1-PP1s that counteracts CDK1 inhibitory effect, allowing the UFB-binding complex to form at the K-chromatin. Simultaneously, this also relieves the suppression of BLM’s centromere unwinding activity. Alternatively, (ii) directly inhibiting CDK1 together with the expression of the MPS1- or PLK1-phosphomutant of BLM can also bypass the centromere protection mechanism.

One of the most important findings of our study is the demonstration of a mitotic kinase network that execute a pathway integral to the control of BTRR/UFB-binding complexes for centromere protection. It is worth noting that both CDK1 and PLK1 have been shown to involve in also protecting telomeres but via different mechanisms^47^. They phosphorylate 53BP1 and RNF8 during mitosis to inhibit their binding to DNA double-stranded breaks on mitotic chromosomes. As a result, the initiation of non-homologous end-joining (NHEJ) is blocked in mitosis. Expression of the corresponding phosphomutants of 53BP1 and RNF8 can bypass the mitotic suppression on NHEJ but cause the fusion of unprotected or damaged telomeres, leading to severe chromosome missegregation^47^. Therefore, our current study reveals the biological significance of programmed mitotic inactivation of chromatin remodellers and DNA repair machineries for the protection of not only the telomeres but also the centromeres.

The centromere protection pathway is governed by several important factors: (1) the integrity of the mitotic BTRR dissolvasome; (2) the interaction between the dissolvasome and PICH; (3) the activity of BLM helicase; and (4) the action of mitotic kinases and protein phosphatases (Fig. 6A). It is generally accepted that the BTRR complexes form stably irrespective of the cell cycle stage^18,22,25,48,49^. Thus, we were surprised to observe its dynamic formation on centromeric chromatin during mitosis. Typically, most protein-protein interaction and co-immunoprecipitation analyses are performed following the elimination of DNA/chromatin, which may overlook its biological influence on protein complex formation. Our live-cell imaging approach thus provides an alternative method to study protein-protein interactions under a physiological condition. An important implication of our findings is that chromatin, and its associated factors, likely exert a significant impact on the stability of DNA-binding complexes. Notably, centromeres and kinetochores are known to harbour numerous protein kinases^50–52^. Potentially, through additional phosphorylation, the high and localised kinase activities may further destabilise the BTRR/UFB-binding complexes when in close vicinity to the centromere. Interestingly, we occasionally observe weak BLM localisation on the protruded pre-UFBs in SGO1-delpeted cells, which are distanced from the core centromere (the zone of high kinase activity). This could be attributed to a partial relief from the phosphorylation-dependent inhibition.

The mitotic phosphorylation has been proposed to stabilise the interaction between PICH and the BTRR complex, as well as to stimulate the DNA unwinding activity on UFBs^21^. However, if this were the case, it would be difficult to reconcile with the findings of Sarlos *et al,* who successfully reconstruct the UFB-binding complex *in vitro* using recombinant proteins that were mostly purified from *E. coli* and yeasts, where specific mitotic phosphorylation is unlikely to be present^8^. More importantly, as demonstrated here, inhibition of mitotic kinases and mutations that abolish BLM mitotic phosphorylation indeed stimulate, rather than compromise, the cellular activity of the UFB-binding complex. In line with this observation, we also show that the RIF1-PP1s activity is required to promote the assembly of the UFB-binding complex induced by PLK1 inhibition, presumably through counteracting the CDK1 inhibitory phosphorylation (Fig. 6B). However, RIF1 is not required for the recruitment of BTRR complex to PICH-bound UFBs during anaphase^9^. Given that the activity of kinases sharply decreases, whereas multiple phosphatases are activated at the metaphase-anaphase transition, it is plausible the inhibitory phosphorylation on the BTRR complex can be also removed by other activated phosphatases. In addition, we show that the mitotic phosphorylation sites on the TOP3A- and RMI1-binding surfaces of BLM can disturb the BTRR complex formation at centromeres. Therefore, our findings collectively support the model that mitotic phosphorylation acts negatively to regulate the UFB-binding factors. Notably, while CDK1 can attenuate stable interaction between BLM and the TRR subcomplex, the phosphorylation sites mapped at the TOP3A/RMI1 binding interfaces of BLM do not align with CDK1 motif consensus. This may suggest the involvement of another kinase(s), downstream of CDK1, in mediating the blocking of the protein-protein interactions. Nevertheless, the characterisation of the BLM and TOP3A interaction also reshapes our understanding on the assembly mechanism of the UFB-binding complex. Although it has been suggested that the BLM helicase and the TRR subcomplex can independently bind to PICH on UFB molecules^8^, our cellular analyses demonstrate that this rarely occurs in anaphase cells. This can be explained by the fact that the binding affinity of PICH for these individual factors is 3 to 5-fold lower than that of the whole BTRR dissolvasome *in vitro*^8^. Thus, multiple PICH-interacting surfaces are probably needed. Interestingly, within the BTRR complex itself, its formation also requires stable interaction of BLM to both TOP3A and RMI1 subunits. Conceivably, these multiple low-affinity interactions, within the BTRR complex and to PICH, likely enable exquisite control of the UFB-binding complex assembly through phosphorylation of different interfaces. This aligns with our hypothesis that regulating the BTRR dissolvasome is one possible means of restraining the UFB-binding complex activity in mitosis.

Disabling CDK1 is sufficient to override the suppression of the BTRR complex to access K-chromatin but inefficient to trigger centromere unwinding. As demonstrated here, this is because of a specific cluster of PLK1-mediated phosphorylation on BLM that limits its centromere unwinding action (Fig. 6B). However, we do not believe that the phosphorylation directly inhibits the enzymatic activity of BLM because phosphorylated BLM isolated from metaphase-arrested cells has been shown to be proficient in DNA unwinding *in vitro*^22^. Intuitively, one might expect a direct allosteric regulation of the helicase domain to be more efficient to suppress BLM, such as altering its ability to bind ATP. However, the critical inhibitory phosphorylation sites identified here reside in the middle of the disordered N-terminal region rather than on the core helicase domain. This may be attributed to higher accessibility of the mitotic kinases to the less structured part of the protein, but how does phosphorylation at the N-terminus of BLM interfere the DNA unwinding at centromere chromatin? One possibility may be through hindering BLM’s accessibility to the DNA substrates and/or co- activators. It is notable that the inhibitory phosphorylation cluster is located between the putative TOPBP1 binding region, close to the essential serine 304 residue for the TOPBP1- BRCT5 domain interaction^28^, and the double helical bundle in the N-terminal (DHBN) domain responsible for BLM dimerization^27,53^(Supplementary Fig. 5D). Mutations in either S304 or the DHBN domain have been shown to compromise BLM anti-crossover but not DNA unwinding activity^18,27^. Thus, it is conceivable that the cluster of phosphorylation might interfere these co-factors, thereby limiting BLM activities at centromeres. Further structural analyses will be required to address these possibilities. In conclusion, we propose that the Bloom syndrome helicase complex can act as a double-edged sword for genome stability. While the active complexes are essential in suppressing genome instability, the lack of temporal regulation can seriously endanger chromosome integrity, particularly at regions of centromere chromatin.

## Supplementary Figure Legends

**Figure S1.**
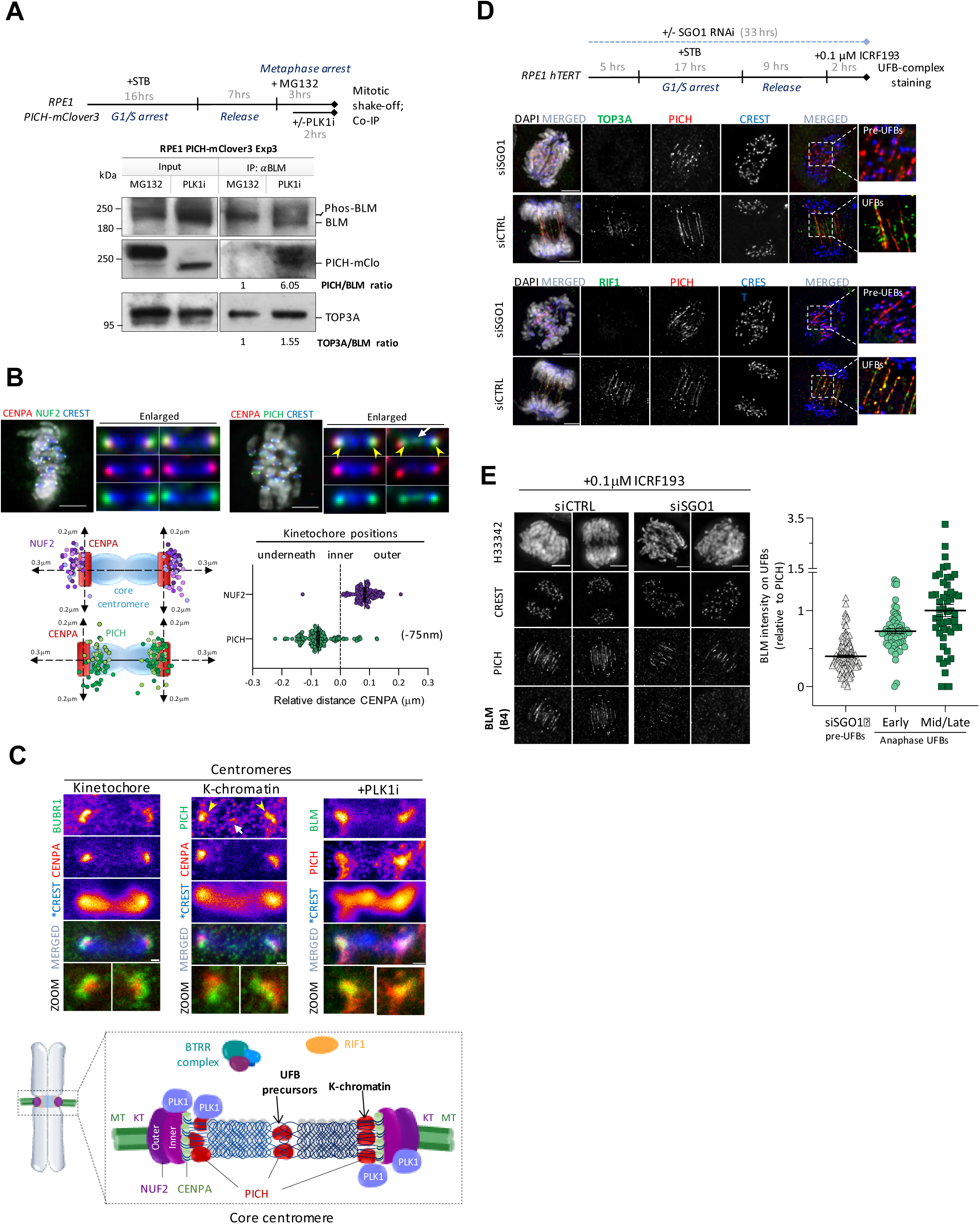
Limited binding of BLM and RIF1 with PICH on pre-UFB structures before anaphase. (A) Repetition of co-immunoprecipitation of BLM, PICH-mClover3 and TOP3A in the presence or absence of PLK1i treatments. Quantification of the normalised ratios are shown. (B) The precise localisation of PICH at human centromeres. Deconvolved images of RPE1 metaphase cells showing the localisation of NUF2 and PICH relative to CENPA. Scale bars, 5 μm. Graphs showing their relative distances to CENPA (114 of NUF2 and 150 of PICH foci were measured; mean ± S.E.M is shown). Arrowheads denote K-chromatin, and the arrow indicates UFB precursors. (C) Super-resolution STED nanoscopy showing the localisation of PICH at K-chromatin (arrowheads) and UFB precursors (arrow), and the colocalisation of BLM after PLK1 inhibition. BUBR1, CENPA and PICH-mClover3 were imaged by STED whereas *CREST by confocal microscopy. Scale bars, 100 nm. A diagram illustrating the K-chromatin and UFB precursors bound by PICH but not the BTRR complex and RIF1. MT, microtubules; KT, Kinetochores; NUF2, Outer kinetochore marker; CENPA, inner kinetochore marker. (D) Representative images showing TOP3A and RIF1 staining on pre-anaphase UFBs and anaphase UFBs in RPE1 cells pre-treated with the indicated siRNA oligos and ICRF193. PICH staining was used to label the UFBs structures and CREST for centromeres. (E) Immunofluorescence staining of endogenous BLM (using mouse B-4 antibody) on pre-anaphase UFBs and anaphase UFBs in RPE1 cells treated with the indicated siRNA oligos. Scale bars, 5 µm. The graph shows the relative intensities (BLM/PICH) on pre-anaphase UFBs and anaphase UFBs. Data is normalised to the average UFB intensity value in mid/late anaphase (total number of UFBs measured per condition: early anaphase UFBs n=143; mid/late anaphase UFBs n= 62, siSGO1 pre-UFBs n=52; mean ± S.E.M is shown).

**Figure S2.**
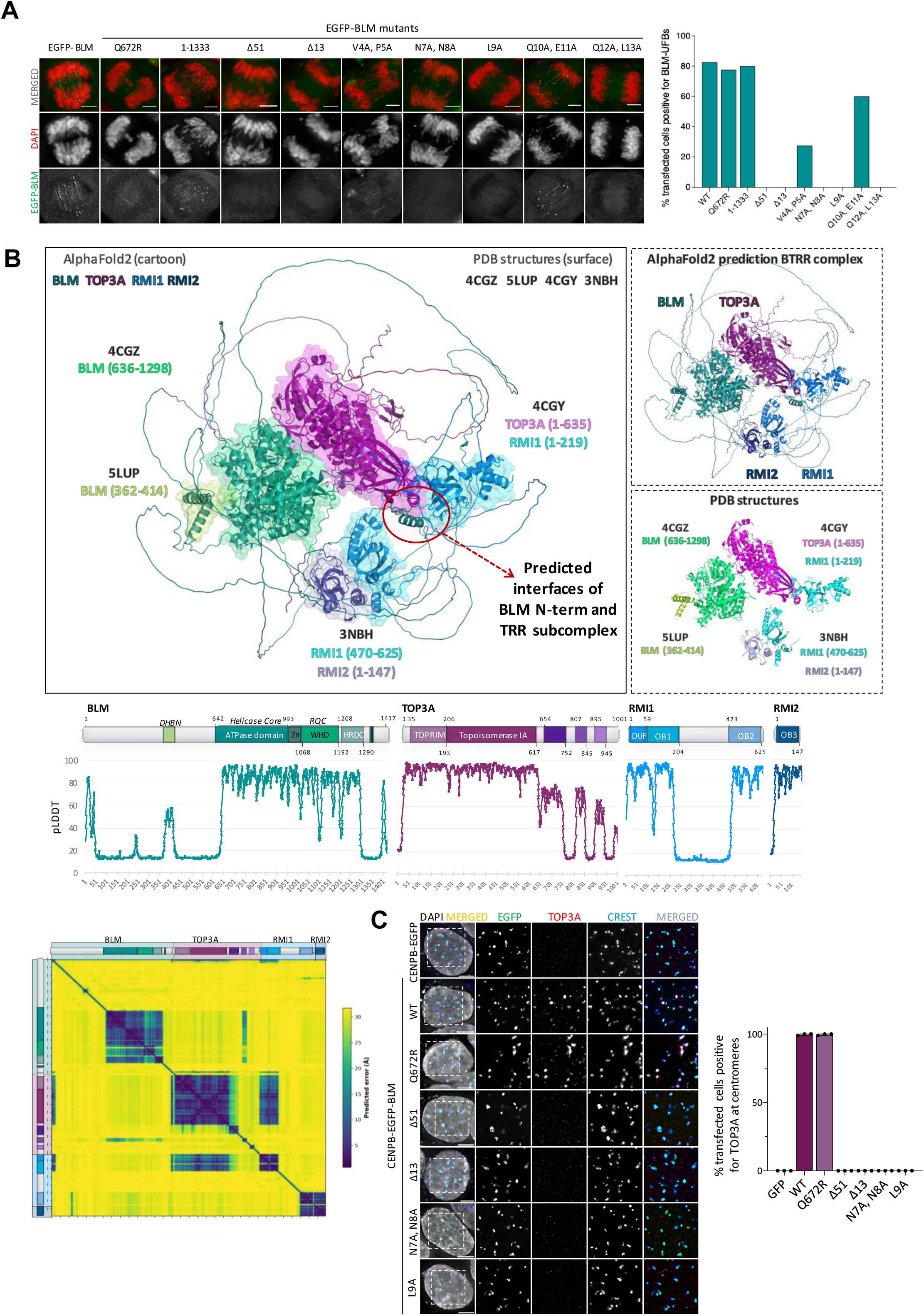
BLM binds to TOP3A and RMI1 using two adjacent N-terminal motifs. (A) HeLa cells were transiently transfected with the indicated EGFP-BLM constructs, synchronised in G2/M by using 9 µM RO-3306 (CDK1 inhibitor), and released in the presence of 1 µM ICRF193 to enrich for anaphase cells containing UFBs. Representative images of BLM UFB localisation. Scale bars measure 5 μm. Graphs showing percentage of transfected anaphase cells showing at least one EGFP-BLM positive UFB. (n, numbers of anaphase cells examined: n=40). (B) Cartoon representations of the BTRR complex predicted by AlphaFold2 and experimental X-ray crystallography structures aligned to the predicted model. PDB:4CGZ, BLM helicase domain; PDB:5LUP, double helical bundle in BLM N-terminus (DHBN); BLM PDB: 4CGY, TOP3A (TOPRIM motif and the ATPase domain) and RMI1 (DUF and OB1-fold domain); PDB: 3NBH, RMI1 (OB2-fold domain) and RMI2 (OB3-fold domain). The red circle highlights the interfaces between the N-terminal region of BLM and the TRR subcomplex (also see Figure 2D). Schematic representation of the subunits of the BTRR complex (DHBN: double helical bundle in N-terminus; RQC: RecQ C-terminal domain; WHD: Winged Helix domain; HRDC: Helicase and RNaseD C-terminal domain; DUF: domain of unknown function; OB: Oligonucleotide/Oligosaccharide-binding domain). Graphs displaying pLDDT scores for each domain within the BTRR dissolvasome. The x-axis represents the residue position within each domain, while the y-axis represents the pLDDT score, indicating the predicted local distance difference between the model and experimental structure. The plot depicts the PAE (Predicted Aligned Error) values for residue-residue alignment within the subunits of the BTRR complex. PAE values are visualised using colour intensity, with dark blue indicating low values and yellow indicating high values. (C) An *in vivo* BLM-TOP3A interaction assay. EGFP-BLM is tethered to centromeres in ΔBLM HAP1 cells by fusing with a truncated CENPB (1-158). The centromeric recruitment of endogenous TOP3A protein was assayed by immunofluorescence staining. Percentages of transfected cells positive for TOP3A at centromeres are shown (total numbers of transfected interphase cells analysed: EGFP n= 176; EGFP-BLM WT n=287; EGFP-BLM Q672R n=254; EGFP-BLM Δ51 n=303; EGFP-BLM Δ13, n=289; EGFP-N7A, N8A n=292; EGFP-L9A n=278; mean ± S.D. is shown, from three independent experiments). Scale bars, 5 µm.

**Figure S3.**
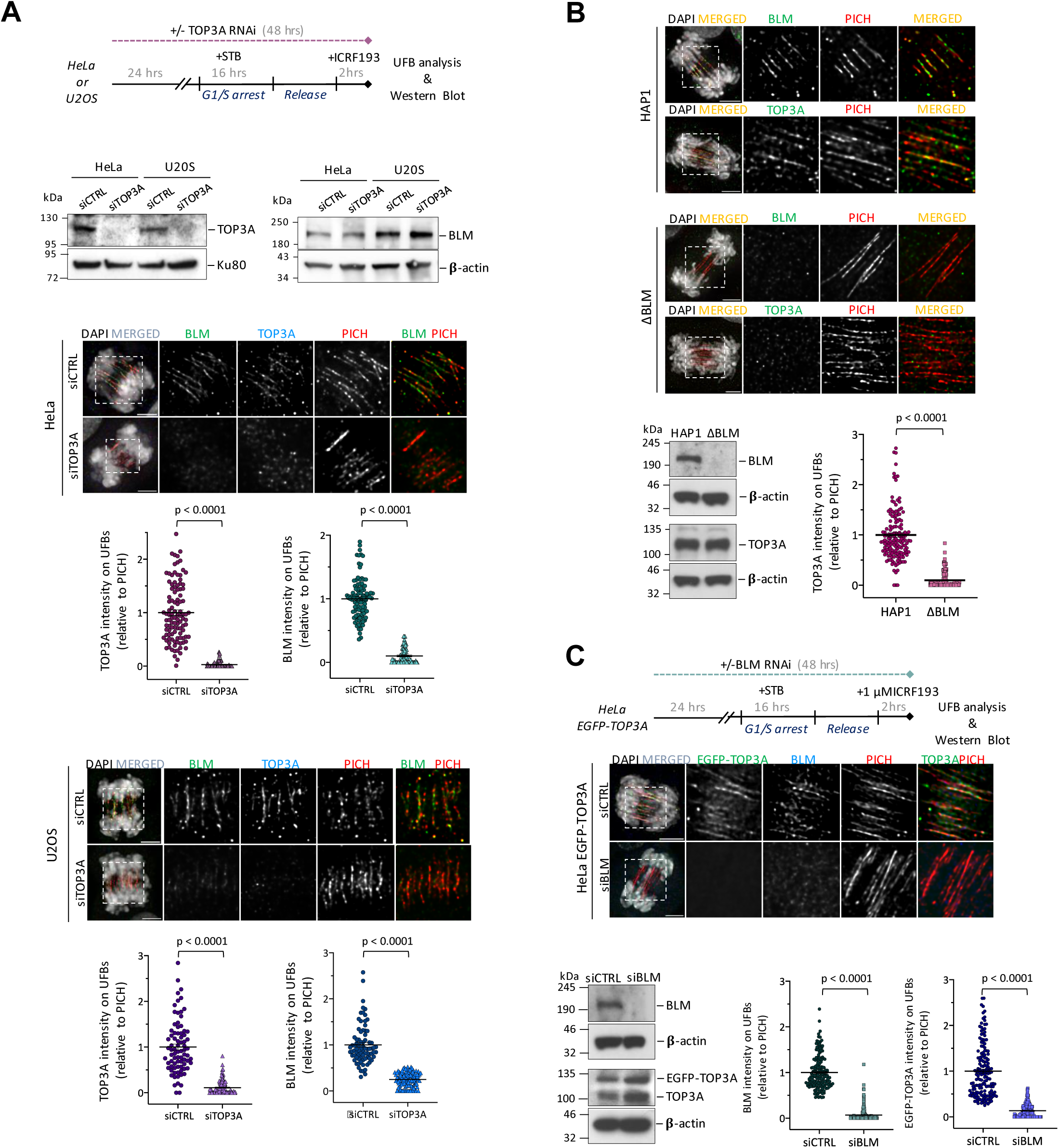
The binding of BLM and TOP3A proteins to anaphase UFBs is interdependent. (A) Experimental outline of TOP3A RNA interference (RNAi) in HeLa, U2OS and RPE1 cells. TOP3A and BLM protein levels were analysed by western blotting after TOP3A knockdown. Representative immunofluorescence images of anaphase HeLa cells (upper panel) or U2OS cells (lower panel) showing BLM, TOP3A and PICH localisation on UFBs after siCTRL or siTOP3A oligos treatments. The graphs show the intensities of BLM or TOP3A relative to PICH on individual UFBs in both conditions. Data are normalised to the average intensity in the control condition (numbers of UFBs measured in HeLa cells; siCTRL n= 105, siTOP3A n= 76 and U20S cells: siCTRL n= 80, siTOP3A n= 114; mean ± S.E.M. is shown). (B) Representative immunofluorescence images of wildtype and ΔBLM HAP1 anaphase cells stained for TOP3A and BLM on UFBs. PICH is used to label the UFBs. BLM and TOP3A protein levels were analysed by Western blotting in both cell lines. The relative TOP3A intensities (to PICH) on PICH-coated UFBs are shown. Data are normalised to the average intensity of TOP3A in wildtype HAP1 cells (numbers of UFBs measured in HAP1 n= 152; in ΔBLM n= 153; mean ± S.E.M. is shown). (C) Experimental outline of BLM RNAi in HeLa cells stably expressing EGFP-TOP3A. Representative immunofluorescence images of anaphase cells showing BLM, TOP3A and PICH localisation to UFBs after siCTRL or siBLM oligos treatments. BLM and EGFP-TOP3A protein levels were analysed by Western Blot. The graphs show quantification of EGFP-TOP3A and BLM intensities relative to PICH. Data are normalised to the average intensity in the control condition (numbers of UFBs measured: siCTRL n= 163; siBLM n= 185; mean ± S.E.M. is shown.). Scale bars, 5 µm.

**Figure S4.**
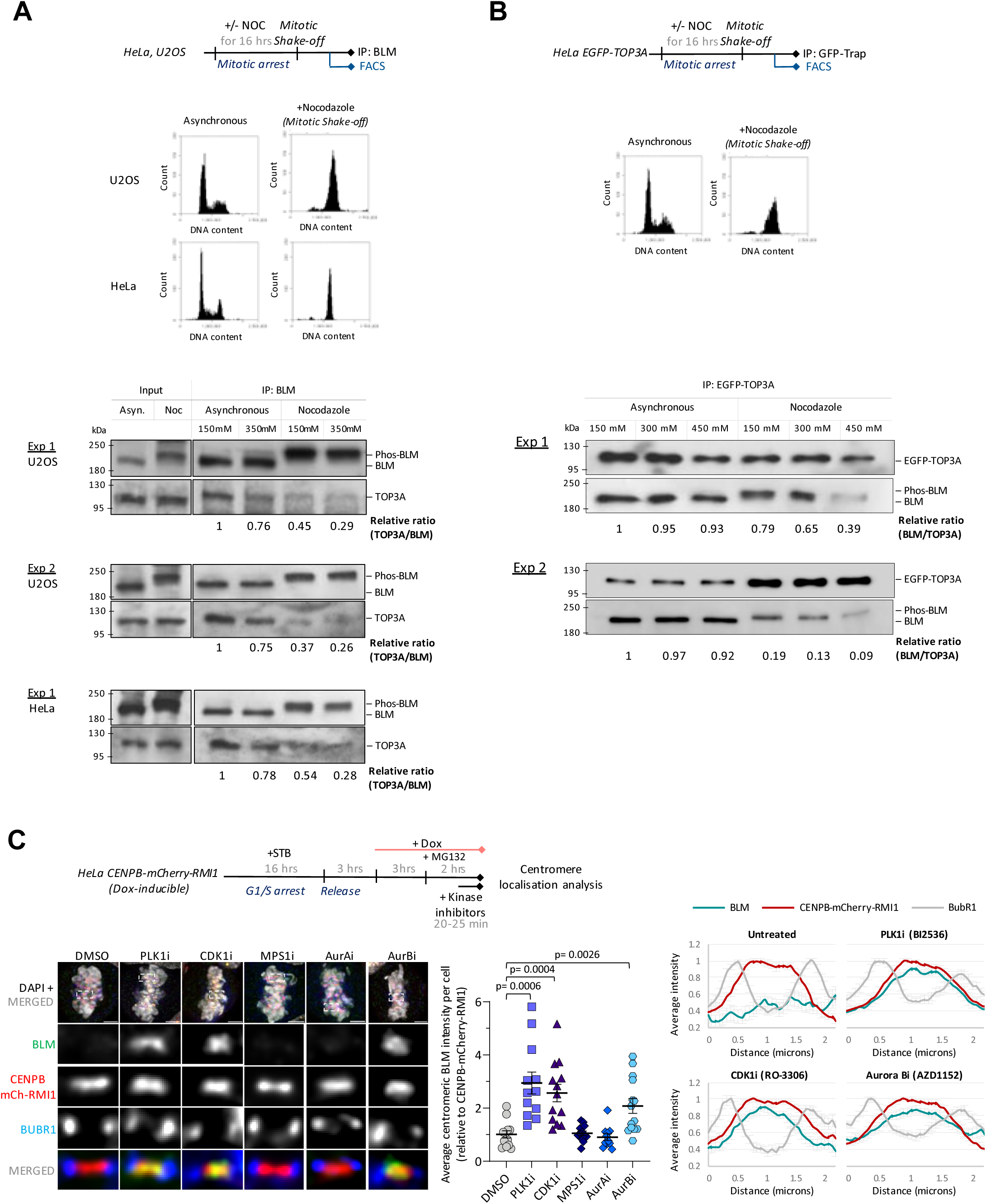
BLM-TOP3A interaction becomes less stable in mitotic cells extracts. (A) Asynchronous and mitotic shake-off cells were collected after the indicated treatments in HeLa and U2OS cells followed by immunoprecipitation of endogenous BLM. FACS (Fluorescent Activated Cell Sorting) profiles of different samples are shown. The immunoprecipitated protein extracts were subjected to washing buffer containing different salt (NaCl) concentrations, as indicated, and analysed by Western Blot. Quantification of the normalised TOP3A/BLM ratio is shown per experiment. (B) Similar to (A) but the EGFP-TOP3A was immunoprecipitated from HeLa cells. Quantification of the normalised BLM/TOP3A ratio is shown per experiment. (C) Experimental outline of centromere localisation analysis under different mitotic kinase inhibition. Representative images of HeLa metaphase-arrested cells stained for endogenous BLM, CENPB-mCherry-RMI1 and the outer kinetochore marker, BubR1. Scale bars, 5 µm. Enlarged images illustrate a single centromere. The graph shows quantification of average centromeric intensity of BLM relative to CENPB-mCherry-RMI1 per cell. Data is normalised to the average intensity in untreated condition (at least 12 cells analysed per condition; numbers of centromere cluster measured: DMSO n=462; PLK1i n= 504; CDK1i n=468; MPS1i n=475; AurAi n=488; AurBi n=549. mean ± S.E.M is shown). Scanline analyses of the centromeres are shown (right panel) (At least 22 centromeres analysed per condition).

**Figure S5.**
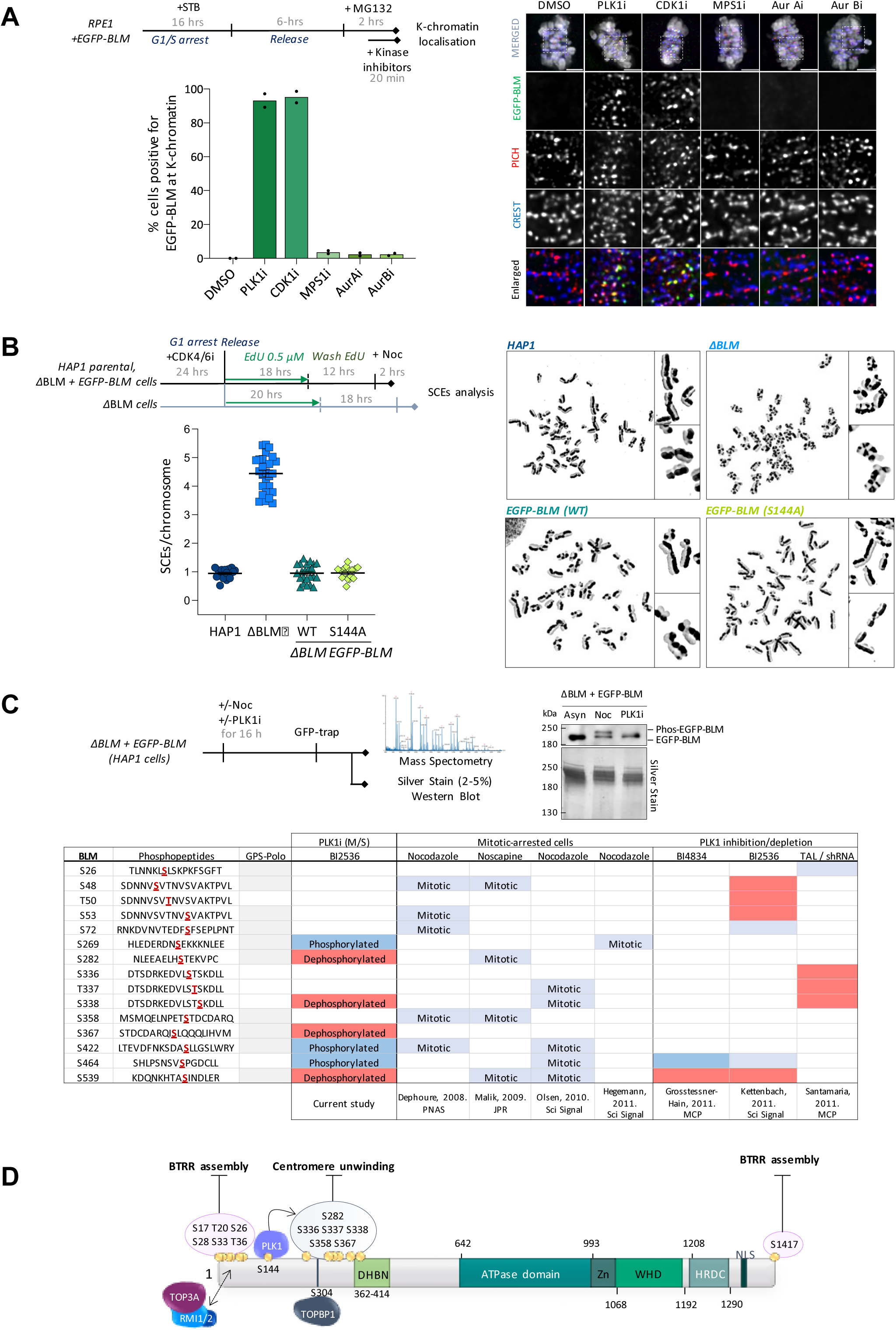
PLK1 and CDK1 prevent the localisation of BLM to K-chromatin in early mitosis. (A) Experimental outline of cell synchronisation for K-chromatin localisation analysis in RPE1 cells stably expressing EGFP-BLM. Representative images of metaphase-arrested cells after treatment with the indicated kinase inhibitors. PICH staining is used to mark the K-chromatin and CREST for centromeres. Quantification of the percentage of cells positive for BLM at K-chromatin (numbers of metaphase cells analysed: DMSO n=126; PLK1i n= 135; CDK1i n=161; MPS1i n=138; AurAi n=130; AurBi n=127; mean is shown from two independent experiments). Scale bars, 5 µm. (B) Experimental outline of cell synchronisation and EdU labelling for SCEs analysis in HAP1, *Δ*BLM and ΔBLM cells complemented with EGFP-BLM WT and S144A mutant. Graph shows quantification of SCE frequency per chromosome, per metaphase spread. Each data point represents the average number of SCEs per chromosome in one spread. (At least 30 metaphases were analysed per condition mean ± S.E.M is shown). Representative images of metaphase spreads of each cell line. (C) Identification of PLK1-dependent phosphorylation in mitotic BLM. Experimental setup for mass spectrometry of the immunoprecipitated BLM proteins from ΔBLM EGFP-BLM cells in the presence or absence of nocodazole/PLK1 inhibitor treatment. Western Blot and silver stain show the enriched EGFP-BLM proteins and the electrophoretic mobility shifts in the indicated conditions. Systematic analysis of PLK1-mediated phosphorylation on BLM: Table shows BLM sites dephosphorylated (sensitive) or phosphorylated (induced) upon PLK1 inhibition (BI2536) identified in the current study. Phosphopeptides sequences are shown, as well as sites predicted to be phosphorylated by Polo-like kinases (PLKs) using bioinformatic software Group-based Prediction System (GPS-Polo). Summary of BLM sites identified in previous proteome-wide analyses (references are depicted) of PLK1-regulated phosphorylation sites in mitotic human cells; either mitotic-specific sites using Nocodazole or Noscapine, or PLK1-dependent phosphorylation sites using shRNA or small-molecule inhibitors of PLK1: BI4834, BI2536 or TAL (ZK-Thiazolidinone). (D) Schematic representation of BLM protein structure, highlighting the cluster of phosphorylation sites identified in this study. Phosphorylation on both the N-terminus and C-terminus abolishes BTRR complex assembly. PLK1-dependent phosphorylation in the middle of the N-terminus of BLM supresses BLM centromere unwinding, downstream of MPS1 phosphorylation on Ser-144 (PLK1-binding site). DHBN domain mediates BLM oligomerisation, while phosphorylation on Ser-304 is required for TOPBP1 binding (BTRR: BLM, TOP3A, RMI1, RMI2; DHBN; double helical bundle in N-terminus; WHD: Winged Helix domain; HRDC: Helicase and RNaseD C-terminal domain; NLS: Nuclear localisation signal).

## Supplementary Movies

**Movie S1.** Centromeric localisation of EGFP-TOP3A (green) in HeLa cells expressing CENPB-mCherry-RMI1 fusion protein (red).

**Movie S2.** Centromeric localisation of EGFP-TOP3A (green) in HeLa cells expressing CENPB-mCherry-BLM(Q672R) fusion proteins (red)

## Methods

### Cell culture and cell lines

Cell lines HeLa, U2OS and HEK293T and their derivates were cultured in DMEM (Sigma) supplemented with 10% foetal calf serum (FCS) (Gibco), 1% L-glutamine and 1% Penicillin/Streptomycin antibiotics. RPE1-hTERT, HAP1 and their derivates were cultured in DMEM/F-12 medium (Sigma) and IMDM (Gibco) respectively, supplemented with 10% FCS (Gibco) and 1% Penicillin/Streptomycin antibiotics. All human parental cell lines used were obtained from the Cell Bank at the Genome Damage and Stability Centre, which were verified by ATCC’s cell line authentication service. All cell lines were periodically tested and confirmed free from mycoplasma (Lonza Mycoplasma testing kit). All cell lines were meticulously maintained at 37°C in a humidified atmosphere containing 5% CO_2_. Stable cell lines in this study were generated by stably transfecting plasmids containing gene of interests either by electroporation using Neon Transfection Kit (ThermoFisher; MPK1025) (RPE1 hTERT cells) at 1350V, for 20ms with 2 pulses or lipid-mediated delivery using FuGENE HD (Promega; TM328) according to the manufacturer’s guidelines. FACS cell sorter (BD FACS Melody) was used to sort and generating stable cell lines expressing the indicated EGFP or mCherry fluorescent tagged proteins. Once recovered the cells were then subjected to antibiotic selection: 800 mg/ml – 1200 mg/ml of G418 for 10 days; 0.25 µg/ml – 0.5 µg/ml of Puromycin for 3 days. Mutations were introduced into plasmids of interest using QuikChange XL Site-Directed Mutagenesis Kit (Agilent Technologies; 200516) according to the manufacturer’s protocol.

### Chemicals and small molecule inhibitors

Doxycycline (Merck; D5207, 0.25-1 µg/ml) was used to induce expression gene of interest using Sleeping Beauty (SB) transposon system (pSBtet vector)^54^. Thymidine (Sigma; T9250, 2 mM), nocodazole (Sigma; SML1665, 50-100 ng/ml), MG132 (Sigma; 474790, 20 μΜ) to control the cell cycle and arrest the cells. Neocarzinostatin (Sigma; N9162, 100 ng/ml) for DNA damage induction. ICRF193 (Merck; I4659, 0.1-1 μΜ) was used to induce UFBs. Mitotic kinase inhibitors: AZD1152 (Sigma-Aldrich; S1147, 100 nM), BI2536 (Cayman Chemical; 17385, 100 nM), MLN8237 (Selleck Chemicals; 1133, 50 nM), Reversine (Axon MedChem; 1629, 1 μΜ), RO-3306 (Sigma-Aldrich; SML0569, 7.5-9 μΜ) and Protein Phosphatase 1 inhibitor: Tautomycetin (Tocris Bioscience; 2305, 1 or 5 μΜ).

### RNA interference

Cells were transfected with siRNA oligonucleotides using Lipofectamine RNAiMAX transfection reagent (Thermo Fisher Scientific; 13778075) following the manufacturer’s guidelines. Non-targeting siRNA pool (Dharmacon ON-TARGET plus Non-targeting Pool—D-001810-10-05. UGGUUUACAUGUCGACUAA; UGGUUUACAUGUUGUGUGA; UGGUUUACAUGUUUUCUGA; UGGUUUACAUGUUUUCCUA). BLM siRNA sequence (Dharmacon ON-TARGET plus Individual—J-007287-08-0005. GGAUGACUCAGAAUGGUUA). TOP3A siRNA sequence (Ambion Thermo Fischer 4392420. CGGCUUGCCUAGUUCUCUA). SGO1 siRNA sequence (Dharmacon ON-TARGET plus SMARTpool—L-015475-00-0005. CAGCCAGCGUGAACUAUAA; GUUACUAUCUCACAUGUCA; AAACGCAGGUCUUUUAUAG; GUGAAGGAUUUACCGCAAA).

### Fluorescence-activated cell sorting (FACS)

Cells were trypsinised, washed with PBS and fixed with 70% ice-cold ethanol, adding dropwise whilst using a vortex. Cells were resuspended in Propidium Iodide (PI) /RNAse staining buffer (9.5 ml 1× PBS, 400 µl 1 mg/ml PI solution, 10 mg/ml RNAase). The resuspended cells were passed through a Falcon cell strainer into a round-bottom tube (Falcon Corning; 352235) for flow cytometry analysis. Cell cycle profiles were then determined and analysed using BD Accuri C6 sampler.

### Immunofluorescence staining

For immunostaining analysis, cells were seeded onto cover glass of No. 1.5 or 1.5H and treated as indicated. Cells were fixed with Triton X-100-PFA buffer (250 mM HEPES pH 7.4, 1× PBS, 0.1% Triton X-100, 4% methanol-free paraformaldehyde) at 4 °C for 20 min, or with PBS–PFA buffer (1× PBS, 3.7 % methanol-free paraformaldehyde) at room temperature for 10 min. Fixed cells washed with PBS and permeabilised with 0.1% of Triton X100 and 0.5 % FCS in PBS for 20min at 4°C, followed by blocking (1× PBS, 0.5 % FCS) at room temperature for 10 min. Cells were incubated with primary antibody at 37°C for 90 min followed by secondary antibody incubation at room temperature for 30 min. Slides were washed with 1× PBS for 5 times at room temperature after antibody incubation. Cells were washed with ultra-pure water and coverslips were airdried before mounting using Vectashield mounting medium, either containing DAPI or without. The latter was used in the cases DNA was stained with Hoechst H33342 (Invitrogen; C10637-G) prior fixation.

Fluorescent images were acquired in a Zeiss AxioObserver Z1 epifluorescence microscopy system equipped with 40x/1.3 oil Plan-Apochromat, 63x/1.4 oil Plan-Apochromat and 100x/1.4 oil Plan-Apochromat objectives and a Hamamatsu ORCA-Flash4.0 LT Plus camera. Z-stack images were acquired at 0.2 μm intervals covering a range from 2–8 μm by using ZEN blue software. Image deconvolution was performed using Huygens Professional deconvolution software with a measured point-spread-function (PSF) generated by 200 nm-diameter TetraSpeck microspheres (ThermoFisher). Classical maximum likelihood estimation method with iterations of 40-60 and a range of 20-60 signal-to-noise was applied. Fiji was used to generate the representative images.

Primary antibody dilution: rabbit BUBR1 (Abcam; ab209998, 1:100), goat anti-BLM (Santa Cruz; sc-sc-7790), mouse anti-BLM (Santa Cruz; sc-365753), rabbit anti-BLM (Abcam; ab2179, 1:100), mouse anti-CENP-A (Abcam; ab13939, 1:100), human anti-CREST (Immuno Vision; HCT-0100, 1:400), alpaca anti-GFP-Atto488 (Chromotek; gba488, 1:200), alpaca anti-GFP-AF488 (Chromotek; gb2AF488, 1:200), rabbit anti-NUF2 (Abcam; ab122962, 1:200), mouse anti-PICH (Abnova; H00054821-BO1P, 1:100), rabbit anti-PICH (Abnova; H00054821-DO1P 1:100), rabbit anti-RIF1 (Bethyl Lab; A300-568A, 1:100), rabbit anti-RIF1 (Bethyl Lab; A300-569A, 1:100), rabbit anti-RPA70 (Abcam; ab79398, 1:200) and rabbit anti-TOP3A (Abcam; ab108493, 1:100); Secondary antibody dilution: donkey anti-goat AF488 (Invitrogen; 1463163, 1:500), donkey anti-mouse AF488 (Invitrogen; A-21202, 1:500), donkey anti-rabbit AF488 (Invitrogen; A-31570, 1:500), donkey anti-mouse AF555 (Invitrogen; A-31570, 1:500), donkey anti-rabbit AF555 (Invitrogen; A-31572, 1:500), donkey anti-mouse AF647 (Invitrogen; A-31571, 1:500), donkey anti-rabbit AF647 (Invitrogen; A-31573, 1:500) and goat anti-human AF650 (Abcam; ab98622, 1:500).

### Time-lapse live-cell microscopy

For live-cell tracking cells were seeded on 2-well or 4-well tissue culture chambers cover glass II (Sarstedt) and were monitored using a Zeiss AxioObserver Z1 epifluorescence microscopy system equipped with a heating and CO_2_ chamber (Digital Pixel). The conditions were kept at 37°C and 5 % CO_2_. Image acquisition was performed using 40×/0.95 Korr Plan-Apochromat (correction collar adjusted to 0.17) or 40×/1.3NA oil Plan-Aprochromat objectives and a Hamamatsu ORCA-Flash4.0 LT Plus camera. ZEN Blue software was used for image and movie acquisition with 3 to 6 z-stacks of 1 μm z-intervals. Images were processed using ImageJ/Fiji software.

### Stimulated Emission Depletion (STED) Super-resolution Nanoscopy

STED images were acquired using the Expert Line Easy3D STED microscope system (Abberior Instruments GmbH). The setup was based on an inverted Olympus IX83 microscope equipped with an objective 100X1.4NA Oil UPLSAPO100XO (Olympus), an automatic XYZ stage, and a real-time autofocusing device. Multichannel confocal images were obtained using excitation lasers of 405 nm, 488 nm, at 561 nm and 640 nm. Stimulated depletion was achieved with a 775 nm STED laser (Abberior Instruments GmbH). Fluorescence signals in predefined channels were detected using avalanche photodiode (APD) detectors. Red and far-red channels STED super-resolution imaging of fixed samples were obtained with typical ∼60 nm XY spatial resolution. Pixel sizes for STED and confocal imaging are 10-15 nm and 100 nm, respectively. The imaging acquisition process was controlled by the Imspector Software (Abberior Instruments Development Team, Imspector Image Acquisition & Analysis Software). H33342 (Invitrogen; C10637 G, 0.25 µg/ml) was used to stain the DNA prior fixation. All primary antibodies were used in a dilution of 1:100. Secondary antibody: goat anti-mouse STAR ORANGE (Abberior; ST580-1001, 1:200), goat anti-rabbit STAR ORANGE (Abberior; ST580-1002, 1:200), goat anti-mouse STAR RED (Abberior, STRED-1001, 1:200), and goat anti-rabbit STAR RED (Abberior, STRED-1002, 1:200). Following immunofluorescence procedure, cells were washed with ultra-pure water and coverslips were airdried before mounting using Mount Liquid Antifade (Abberior). The excitation and depletion laser powers for STAR RED are 3-5% and 25-45%, respectively. The excitation and depletion laser powers for STAR ORANGE are 15-25 and 25-35%, respectively. Line accumulation is set between 15 to 20. Images were further processed using ImageJ/Fiji software.

### Imaging analysis

CENPA-PICH/CENPA-NUF2 coordinates: The measurements of the distance between foci in two channels of an image were done by using Spot Pair Distance Tool in FIJI software. The tool searches within a focus/box radius, typically± 5px, for a local maximum in the two pre-selected analysis channels. The centre-of-mass around each maximum, typically± 2px, is computed as the centre of intensity for each channel. Dragging from the clicked point creates a reference direction. The Euclidean distance between the centres is reported, optionally with the signed XY distance and angle relative to the reference direction. Visual guides are overlaid on the image to assist in spot selection and direction orientation. Available in the latest GDSC (Genome Damage and Stability Centre) Fiji plugins.

UFB intensity measurements: UFBs were stained for the UFB-binding complex components, including BLM, TOP3A and PICH following immunofluorescence protocol. FIJI software was used to determine the intensity of individual UFBs by drawing a scanline along the entire length of each UFB thread using PICH channel as a reference, which allowed the measurement of the absolute intensity of other channels corresponding to either BLM or TOP3A. The intensity of different channels was subjected to background correction prior to further analysis.

Centromere localisation analysis: The intensity of each centromere/cluster was determined using the Find Foci GUI Tool in FIJI software, which locates all the points of maximum intensity in the centromeric regions. This was used to generate a mask image containing all the pixels in the peak region above background in the CENPB-mCherry or PICH channel by using the Otsu thresholding method. The intensities of different channels within the centromeric masked regions were then measured using the Mask Analyser Channel Tool available in the latest GDSC (Genome Damage and Stability Centre) Fiji plugins.

### Western Blot

Cells were trypsinised and washed with ice-cold PBS once followed by protein extraction in Lysis Buffer A or B (Lysis buffer A: 50 mM Tris-HCl pH7.5, 300 mM NaCl, 5mM EDTA, 1% Triton X-100, 1 mM DTT, 1 mM PMSF; Lysis buffer B: 50 mM Tris-HCl pH7.5, 150 mM NaCl, 1% Triton X-100, 1.25 mM DTT and 1 mM PMSF, 25U/ml benzonase (Sigma-Aldrich; E1014). The lysis buffers were supplemented with Protease inhibitor cocktail cOmplete protease inhibitor (Roche; 11697498001) 1 tablet/50 ml and when required, phosphatase inhibitor PhosSTOP (Roche; 4906845001) 1 tablet/10 ml. Cell lysate was incubated on ice for 15-20 mins and centrifuged at 15000 rpm for 20 min at 4°C. Protein concentration was quantified using a Bradford assay (Bio-Rad). Samples were separated by SDS-PAGE, transfer onto Amersham Hybond PVDF membranes, 0.2 □m (GE Healthcare Lifesciences, 10600021) and blotted with the indicated antibodies following standard procedures.

Primary antibodies: mouse anti-β-actin (Sigma; A5441, 1:5000), goat anti-BLM (Bethyl Lab; A300-120, 1:2000), rabbit anti-BLM (Abcam; ab2170, 1:500), rabbit anti-GFP (Abcam; ab290, 1:2000), rat anti-GFP (Chromotek; 3h9-100, 1:2000), rabbit anti-Ku80 (Abcam; ab80592, 1:5000), mouse anti-PICH (Abnova; H00054821-BO1P, 1:500), rabbit anti-RIF1 (Bethyl Lab; A300-568A, 1:1000), rabbit anti-RIF1 (Bethyl Lab; A300-569A, 1:1000), rabbit anti-TOP3A (Proteintech; 14525, 1:500). HRP-conjugated secondary antibodies: rabbit anti-goat (Agilent; P0160, 1:60000), goat anti-mouse (Abcam; ab6780, 1:25000), donkey anti-rabbit (ECL/Sigma; NA9340, 1:40000), and goat anti-rat (ECL/Sigma; GENA935, 1:25000).

### Co-immunoprecipitation

HEK293T cells were transiently transfected with EGFP-BLM or EGFP-TOP3A plasmids or the □BLM HAP1 cells with EGFP-TOP3A plasmids. Additionally, HeLa cells stably expressing EGFP-TOP3A were also used to perform mitotic analysis of protein-protein interaction. The cell pellet was lysed in the lysis buffer A or B as the immunoblot procedure. Input samples were collected prior to standard immunoprecipitation procedure. The lysate was applied on the equilibrated GFP-Trap_Magnetic Agarose beads (GFP-Trap_MA) or GFP-Trap_Dynabeads (GFP-Trap_M-270) and rotated for 1 hour at 4°C. Then the samples were collected by a magnet and washed three-five times in the diluting buffer (50 mM Tris-HCl pH7.5, 150 mM NaCl, 0.5 mM EDTA). Concentrations of NaCl varied ranging from 150 mM to 450 mM when indicated. The proteins bound to the beads were resuspended in 1X Laemmli Sample Buffer (BioRad) with 5% □-mercaptoethanol (Sigma, M6250) and boiled for 5 minutes. Supernatant was analysed by western blot.

Endogenous BLM was pull downed in HeLa, U2OS or RPE1 PICH-mAID-mClover3 cells using rabbit anti-BLM (Abcam; ab2179). Cells were lysed in lysis buffer B containing benzonase. Input samples were collected prior to standard immunoprecipitation procedure. Following a pre-incubation step of the cell lysates with the antibody for 1 hour at 4 °C, the mixture was applied to DynaGreen^TM^ Protein A/G magnetic beads (Invitrogen; 80104G) and rotated for 1 hour at 4 °C. Then the samples were washed three times in the diluting buffer (50 mM Tris-HCl pH7.5, 150 mM NaCl, 0.5 mM EDTA). Various concentrations of NaCl were applied during the washing step as indicated in the individual experiments. The proteins bound to the beads were resuspended in 1X Laemmli Sample Buffer (BioRad) with 5% □-mercaptoethanol (Sigma; M6250) and boiled for 5 minutes. Supernatant was analysed by western blot.

### Mass Spectrometry

HAP1 ΔBLM stably expressing EGFP-BLM wildtype was used for mass spectrometry analysis following immunoprecipitation procedures using GFP-Trap assay. The cell pellet was lysed in lysis buffer B (as indicated in ‘Western Blot’ methods section). The lysate was applied on the equilibrated GFP-Trap_Magnetic Agarose beads (GFP-Trap_MA) and rotated for 1 hour at 4°C. Then the samples were collected by a magnet and washed three-five times in the diluting buffer (50 mM Tris HCl ph 7.5, 150 mM NaCl, 0.5 mM EDTA). The proteins bound to the beads were resuspended in 1X Laemmli Sample Buffer (BioRad) with 5% □-mercaptoethanol (Sigma; M6250) and boiled for 5 minutes. Supernatant was run in an SDS-PAGE gel, which was stained by Coomassie blue. Bands corresponding to the EGFP-BLM size were extracted and frozen at -80 °C. Liquid Chromatography with tandem mass spectrometry (LC-MS) experiment was performed by the BSRC Mass spectrometry and Proteomics Facility at the University of St. Andrews. In brief, gel bands were reduced, alkylated and digest followed phospho-peptide enrichment on TiO2 (Titanium dioxide) columns. The eluted phospho-peptides were subsequently analysed on their LC-ESI-MS instrument. The data was searched against the NCBI (National Center for Biotechnology Information) database containing all species and analysed using the Mascot Server. In a second run of mass spectrometry the immunoprecipitation of EGFP-BLM wildtype was carried out in a similar manner. A fraction of the beads was resuspended in 1X Laemmli Sample Buffer (BioRad) with 5% □-mercaptoethanol (Sigma, M6250) and boiled for 5 minutes. This was analysed by silver staining, using Silver Staining Kit (Pierce) and Western Blot. After the washing steps, beads were additionally washed tree times with 1× PBS, to remove detergents and free amines, and frozen at -80°C. Trypsin digestion and TMT-3plex (Tandem mass tag) mass spectrometry was performed at Institute of Cancer Research (ICR). In brief, samples were solubilised, reduced, alkylated, and digested followed by labelling with the TMTpro multiplexing reagents (Thermo Scientific). Prior to LC-MS analysis of TMT-labelled peptides were fractionated with high-pH reversed-phase (RP) chromatography and enriched with immobilised metal ion affinity chromatography. The SequestHT search engine was used to analyse the data.

### AlphaFold2 protein structure prediction

For modelling of the BTRR dissolvasome, FASTA protein sequences were submitted to AlphaFold version 2.3.2^29^ run on Apocrita, the Queen Mary, University of London High Performance Cluster. For both complexes, 5 models with 5 seeds were produced using multimer mode and the top ranked model was relaxed and taken for figure production using PYMOL Version 2.2.2 (The PyMOL Molecular Graphics System, Version 2.2.2 Schrödinger, LLC) to produce structural figures and AlphaPickle Version 1.5.4 (M. J. Arnold, 2021) for statistical plots.

### Metaphase spreads preparation

Cells were first synchronised using single thymidine block and released to S phase for the indicated period. Two hours before the metaphase spread preparation, the cells were treated with 60 nM of the PLK1 inhibitor, BI2536 (Cayman Chemical; 17385). Mitotic cells were collected by mitotic shake-off. Cells pellets were resuspended and mixed very gently in 10 ml of pre-warmed hypertonic solution (0.075 M KCl) followed by an incubation at 37 °C for 10 min. The swollen cells were centrifuged, fixed, and washed twice with the Carnoy’s Fixative (3:1; Methanol: Acetic Acid) according to the standard chromosome spread preparation. Chromosome spreads were dropped onto glass slides and stored at room temperature prior to FISH hybridisation.

### Centromere and Telomere Fluorescence *in Situ* Hybridisation (FISH)

The Peptide nucleic acid (PNA) probe hybridisation of chromosome spreads followed the manufacturers guidelines (DAKO Agilent & PNAbio). Chromosome spreads were fixed in 3.7% PFA and washed using a gradient ice-cold ethanol wash series 70%, 90% and 100% EtOH. Slides were then air-dried prior to PNA probe addition to the spreads: centromere, CENTB-FAM (PNAbio; F3006) and Telomere TelG-Cy3 (PNA FISH kit, Dako Agilent; K532611-8 or PNAbio; F1006). A coverslip (18x18mm) was added to the probe and the slide was co-denatured at 80 °C for 30 sec to 1 min and hybridized at room temperature in a humidifying chamber for 2 hours. Slides were then washed in Wash Solution (Dako Agilent) at 65°C and dehydrated again in the ice-cold ethanol series. Slides were air dried and counterstained using Vectashield with DAPI.

### Sister chromatid exchange (SCE) assays

Cells were arrested in G1 using CDK4/6 inhibitor (Palbociclib, Merck; PZ0383, 1 µM) and release into S phase in the prescence of EdU (Invitrogen; C10637, 0.5 µM) for 18-22 hours. EdU was washed from the cells, which were kept in fresh medium for 12-18 hours. To trap the mitotic cells, cells were arrested using Nocodazole (Sigma; SML1665, 50 ng/ml) for 2 hours prior to metaphase spread preparation. EdU was detected using Click-iT Plus EdU labelling kits (Alexa Fluor 488, Invitrogen; C10637). Slides were air dried and counterstained using Vectashield with DAPI.

### Quantification and statistical analysis

Statistics analysis was performed by using GraphPad Prism software version 9 by two-tailed unpaired Student’s t-test. Data were presented as the mean ± standard deviation (s.d.) unless specified. Probability value ‘p ≤ 0.05’ was considered to be significant.

## Acknowledgments

We would like to thank Professor Marcel van Vugt (University of Groningen) for the ΔBLM HAP1 cells, Professor Steve Jackson (University of Cambridge) for the ΔRIF1 RPE1 cell line, Professor Anne Donaldson and Dr. Shin-Ichiro Hiraga (University of Aberdeen) for the plasmids of EGFP-tagged human RIF1 WT and PP1bs, Professor Mark Burkard for the cDNA of CENPB (residue 1-158), and Dr Gary Ying-Wai Chan (the University of Hong Kong) for the CRISPR plasmids for tagging endogenous PICH with mAID-mClover3. We would like to acknowledge the BSRC Mass Spectrometry and Proteomics Facility (University of St. Andrews) and the Proteomics Core Facility (The Institute of Cancer Research, ICR) for the mass spectrometry services. This research utilised Queen Mary’s Apocrita HPC facility, supported by QMUL (Queen Mary University of London) Research-IT. We thank Professors Aidan Doherty, Evi Soutoglou, Timothy Humphrey and the members of Chan Lab for their valuable comments on this manuscript. Thanks also go to Dr. Anthony Oliver for gifting the PYMOL software license (University of Sussex). We thank Jonathan Wing for the help with cell sorting (University of Sussex), and Dr. Yan Gu and the Wolfson Centre for Biological Imaging (University of Sussex) for providing excellent microscopy and imaging facilities. The current research was supported by the Sir Henry Dale Fellowship (104178/Z/14/A) provided by Wellcome Trust and the Royal Society, and Wellcome Trust Career Development Award (225348/Z/22/Z) to K.L.C.

## Author Contributions

M.F.C, and K.L.C. contributed to the conceptualisation and development of the project. M.F.C., E.K., T.O., U.A. and K.L.C. performed the experiments, collected, and analysed the data. A.H. developed the Fiji imaging analysis plugins. M.D. performed the AlphaFold2 structural predication and analysis. M.F.C, and K.L.C. prepared figures and wrote the manuscript.

## Data availability

All data are provided in the supplementary information. Microscope images and other data that support the conclusions of this manuscript are available upon request.

